# *Caenorhabditis elegans* ADAR editing and the ERI-6/7/MOV10 RNAi pathway silence endogenous viral elements and LTR retrotransposons

**DOI:** 10.1101/825315

**Authors:** Sylvia E. J. Fischer, Gary Ruvkun

## Abstract

Endogenous retroviruses and LTR retrotransposons are mobile genetic elements that are closely related to retroviruses. Desilenced endogenous retroviruses are associated with human autoimmune disorders and neurodegenerative diseases. *C. elegans* and related *Caenorhabdites* contain LTR retrotransposons and, as described here, numerous integrated viral genes including viral envelope genes that are part of LTR retrotransposons. We found that both LTR retrotransposons and endogenous viral elements are silenced by ADARs (*a*denosine *d*eaminases *a*cting on *d*ouble-stranded RNA (dsRNA)) together with the endogenous RNAi factor ERI-6/7, a homolog of Mov10 helicase, a retrotransposon and retrovirus restriction factor in human. siRNAs corresponding to integrated viral genes and LTR retrotransposons, but not to DNA transposons, are dependent on the ADARs and ERI-6/7; on the contrary, siRNAs corresponding to palindromic repeats are increased in *adar-eri* mutants because of an antiviral RNAi response to dsRNA. Silencing of LTR retrotransposons is dependent on downstream RNAi factors and P granule components but is independent of the viral sensor DRH-1/RIG-I and the nuclear Argonaute NRDE-3. The activation of retrotransposons in the ADAR- and ERI-6/7/MOV10-defective mutant is associated with the induction of the Unfolded Protein Response (UPR), a common response to viral infection. The overlap between genes induced upon viral infection and infection with intracellular pathogens, and genes co-expressed with retrotransposons, suggests that there is a common response to different types of foreign elements that includes a response to proteotoxicity presumably caused by the burden of replicating pathogens and expressed retrotransposons.

**SIGNIFICANCE:** Silencing of transposable elements and viruses is critical for the maintenance of genome integrity, cellular homeostasis and organismal health. Here we describe multiple factors that control different types of transposable elements, providing insight into how they are regulated. We also identify stress response pathways that are triggered upon mis-regulation of these transposable elements. The conservation of these factors and pathways in human suggests that our studies in *C. elegans* can provide general insight into the regulation of and response to transposable elements and viruses.

## INTRODUCTION

Transposons can have profound effects on cellular function such as disruption of gene function by transposon insertion, cell death as a consequence of transposon-induced double stranded breaks, and genomic rearrangements caused by homologous recombination between repeat elements. Retrotransposons and endogenous retroviruses may affect cellular function through overexpression of toxic proteins or the activity of retroviral proteins on endogenous sequences. Palindromic repeat elements such as those formed by adjacent but inverted insertion of vertebrate Alu elements into genes can cause the accumulation of dsRNA.

Many RNA viruses replicate via a dsRNA intermediate, which is detected to trigger an interferon-based antiviral response in mammals and an RNA interference response in invertebrates. To avoid an anti-viral response to endogenous palindromic dsRNAs when in fact there is no viral infection, ADARs edit adenosines to inosines in endogenous dsRNA destabilizing the RNA duplex and thus preventing recognition by RIG-I and MDA-5/IFIH1 (1) (2). The importance of ADAR editing is underscored by the severe defects caused by ADAR mutations in human and mouse. Human ADAR1 mutations can cause Aicardi-Goutières Syndrome, which manifests as a congenital viral infection (3). Similarly, a severe form of age-related macular degeneration, geographic atrophy, is linked to the accumulation of Alu dsRNA and an inflammosome response, thought to be caused by the loss of Dicer1 activity with age (4, 5).

The nematode *C. elegans* has, in addition to the *adr-1* and *adr-2* ADAR genes, a very active and diversified RNAi machinery that can act on dsRNA; ADAR activity competes with Dicer for dsRNA (6). *C. elegans* RNAi pathways regulate gene expression and silence DNA transposons through piRNAs and endogenous siRNAs. The *C. elegans* homolog of the human Mov10 helicase, ERI-6/7, is an endogenous RNAi factor that acts in an RNAi pathway that regulates the expression of about one hundred *C. elegans* genes of probable viral origin (7). *C. elegans* defective in ADAR editing (the *adr-1; adr-*2 double mutant) or defective for the ERI-6/7/helicase RNAi pathway are healthy. However, *C. elegans* defective in both the ADAR and the ERI-6/7 pathways are synthetically very sickly (see below and (8)). We used the *adar*- and *eri*-*6/7*-defective mutant to genetically and transcriptionally study the interactions between the ADAR editing and endogenous RNAi. Similar to the rescue of lethality caused by the absence of ADARs by loss of MAVS (a factor downstream of RIG-I) in mammals (9), the morphological defects of the *adar*- and *eri*-*6/7*-defective mutant are rescued by loss of *drh-1*, a gene encoding the *C. elegans* ortholog of the viral sensor protein RIG-I. Loss of ADAR editing in enhanced RNAi mutants results in anti-viral RNAi response to dsRNA from palindromic repeat elements that are normally edited by ADARs but are now a substrate of the RNAi machinery, resulting in an accumulation of novel siRNAs and accompanied by an upregulation of RNAi machinery.

In addition, we show that LTR retrotransposons are regulated by ADARs and ERI-6/7/MOV10. The human ERI-6/7 homolog Mov10 is an antiviral factor and restricts retrotransposon activity (10, 11). Mov10 binds to 3’UTRs and is thought to clear these of 3’UTR binding proteins and/or secondary structures (12); Mov10 also has helicase-independent anti-viral activity(13). *C. elegans* harbors fragments and full-length copies of at least 20 families of LTR-containing retrotransposons that belong to the Ty3/gypsy and the BEL/Pao classes (14, 15). LTR retrotransposons and endogenous retroviruses (ERVs) are related to retroviruses from which they differ in the absence of an envelope protein gene or the presence of inactivating mutations. We found that retrotransposons (but not DNA transposons) are silenced through a mechanism that requires ADARs and ERI-6/7 for siRNA generation. The ERI-6/7 helicase, together with the Argonaute ERGO-1, acts in an RNAi pathway that silences genes that are likely to be remnants of viruses integrated in the genome (7). We here show that the ERI-6/7 helicase regulates expression of viral envelope proteins that may have been acquired by LTR retrotransposons, suggesting that these elements potentially are endogenous retroviruses. Viruses, and also retrotransposons, co-opt the ER for maturation and assembly (16, 17). The strong induction of the UPR in *C. elegans* defective in ADAR editing and with a defective ERI-6/7 endogenous RNAi pathway is likely a consequence of ER stress caused by overexpression of retrotransposons and viral proteins: we found a tight co-expression of retrotransposon genes and UPR genes, under conditions such as loss of silencing factors, loss of germ line P granules and in aged animals. The similarities between LTR retrotransposons that have integrated into the genome and viruses that invade the cell extends beyond sequence similarities to the silencing mechanisms of these elements and to the transcriptional response to these elements.

## RESULTS

### ADAR editing enzymes and the ERI-6/7 RNAi pathway interact to prevent the induction of a toxic anti-viral response to endogenous dsRNA

ADAR editing is essential in mammals, as demonstrated by the lethality of ADAR1 and ADAR2 mutants in the mouse, and the severe defects caused by hypomorphic mutations in human ADAR1. In contrast, the two *C. elegans* ADAR genes (Figure 1A) are dispensable for viability. Because both the ADARs and RNA interference (RNAi) can act on dsRNA, and both have anti-viral activity in mammals, we tested for synthetic interactions between *adar* null mutants and RNAi pathway mutants. We found that *C. elegans* mutants that are defective in the two ADAR genes and defective in the ERI-6/7 helicase (a homolog of human MOV10 and MOV10L1; Figure 1A) that acts in an RNAi pathway that targets recently acquired genes, are severely compromised (for simplicity we call the *adr-1 eri-6/7; adr-2* triple mutant ‘*adar-eri’* from here on): the animals display defects in vulva morphology, causing frequent rupture (Supplemental Figure 1A), appear starved and are partially lethal. As a result, they produce almost no offspring (Figure 1B). The morphological phenotype was reproduced in worms deficient in ADAR (with different *adr-1* and *adr-2* mutations) in combination with a number of mutations in genes acting in the same pathway as *eri-6/7*, including the Argonaute gene *ergo-1* (data not shown). Synthetic lethality of the *adr-1; adr-2* mutations with the RdRP *rrf-3* was reported by Reich et al. (8).

**Figure 1.**
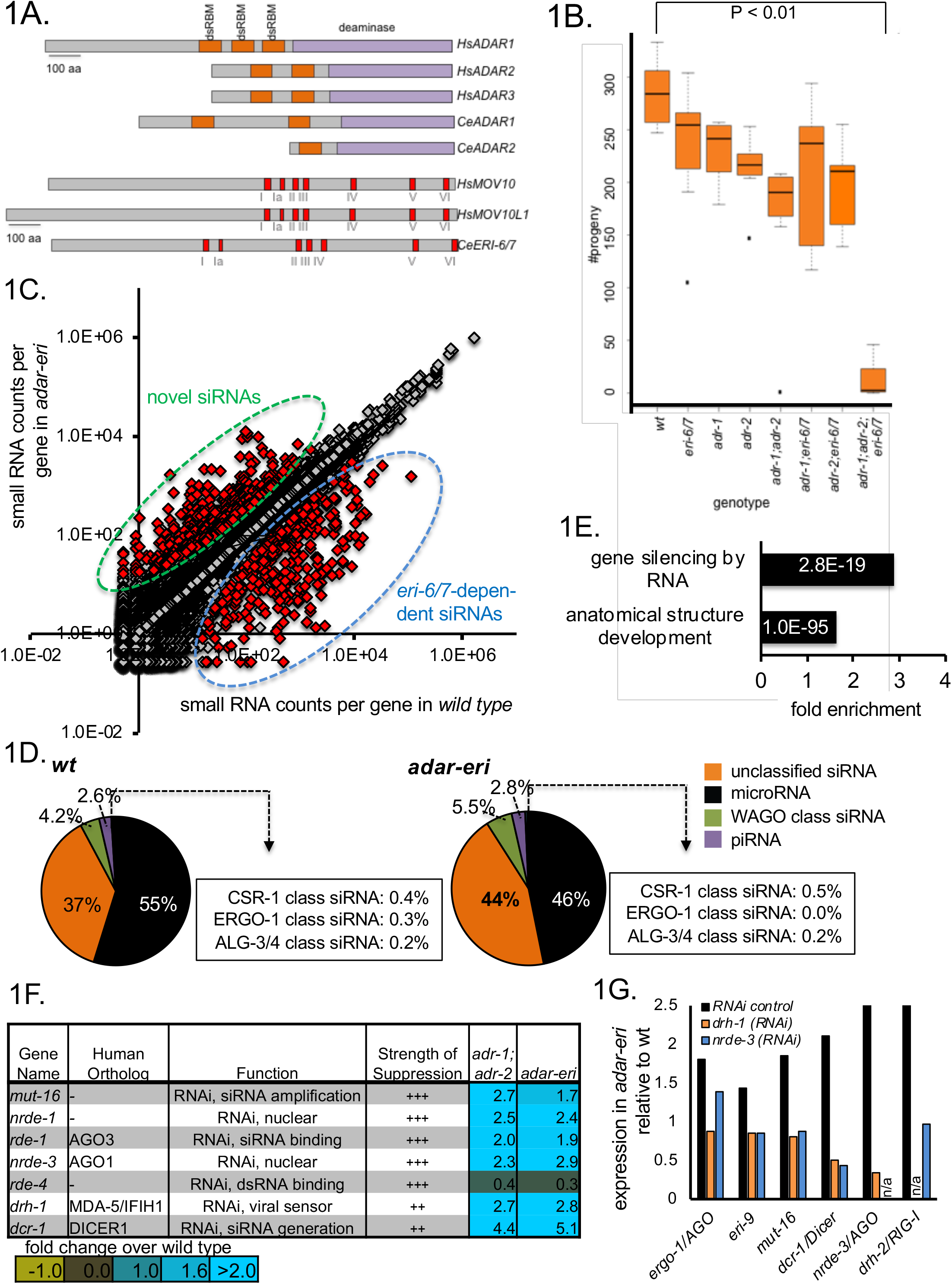
A. ADAR and MOV10 proteins encoded in the human and *C. elegans* genomes. Indicated are double-stranded RNA-binding motifs (in orange), the deaminase domains (in purple), and the Superfamily I sequence motifs that coordinate the NTP and nucleic acid binding (in red). B. *adar-eri* triple mutants have a severely reduced fecundity, whereas *adr-1*, *adr-2* and *eri-6/7* single and double mutants show only modest reductions in brood size. C. A subset of genes produces more endogenous siRNAs in a*dar-eri* triple mutants than in wild type. The number of siRNAs per gene in *adar-eri* is plotted against the number of siRNAs per gene in wild type. D. *adar-eri* mutants produce novel siRNAs that do not act in canonical endogenous siRNA pathways. siRNAs are classified by the specific Argonaute protein they are known to bind to or depend on. E. GO-term enrichment analysis of genes upregulated in *adar-eri* (and *adar*) mutants identifies RNA silencing. F. Expression levels of suppressors of *adar-eri* phenotypes relative to wild type. Differential expression for all genes shown is significant. G. Expression of RNA silencing genes in *adar-eri* triple mutants after knockdown of *drh-1/RIG-1, nrde-3/AGO* or control RNAi.

The *eri-6/*7 single mutant is defective in RNAi-mediated silencing of recently acquired genes. To identify the cause of the near-inviability of the *adar-eri* triple mutant, we sought to determine whether *adar-eri* triple mutants have additional defects in RNAi pathways. We generated sequencing libraries of small RNA between 18 and 28 nucleotides in length that includes microRNA, piRNA and endogenous siRNAs (small RNA mapping to genes, transposons etc). Whereas the *eri-6/7* and *adr-1;adr-2* mutants have grossly similar levels of siRNAs versus microRNAs, the *adar-eri* triple mutant has an increase in siRNAs (Figure 1C). These siRNAs are 22 nucleotides long and start with a 5’G, typical of siRNAs produced by RNA-dependent RNA polymerases (Supplemental Figure 1C).

The endogenous siRNAs were mapped to genes, pseudogenes and transposons that are regulated by particular Argonautes. *adar-eri* triple mutants display the expected loss of ERGO-1 Argonaute class siRNAs that are dependent on ERI-6/7 (Figure 1D); these siRNAs are also missing in the *eri-6/7* single mutant (Supplemental Figure 1D). However, the largest change is an increase in siRNAs corresponding to loci that are not known to be regulated by the ERI-6/7 pathway, or to mostly coding loci that do not produce siRNAs in wild type (Figures 1C and 1D). A previously identified class of siRNAs present in *adar* mutants is not further increased in *adar-eri* triple mutants (Supplemental Figure 1E). Over 450 genes show an increase in siRNAs of 3-fold or more in the *adar-eri* triple mutant, of which 40% are increased only in the *adar-eri* triple mutant (Supplemental Figure 1F).

The increase in siRNAs suggests that *adar-eri* triple mutants activate RNAi too intensely. This hypothesis is supported by transcriptome analysis of the *adar*, *eri* and *adar-eri* triple mutants. We found that genes with GO term ‘silencing by RNA’ are significantly enriched (P value = 2.8E-19) among genes with increased expression in the *adar* mutant and in the *adar-eri* triple mutant versus wild type (Figure 1E). A survey of known RNAi factors indeed shows a 4- to 14-fold induction of many of the core RNAi factors in both *adar* and *adar-eri* triple mutants, but not in the *eri-6/7* mutant (Supplemental Figure 1G, Figure 1F). Also, gene inactivations of RNAi factors including DRH-1/RIG-1, RDE-1/AGO, and NRDE-3/AGO suppress the lethality of the *adar-eri* triple mutant (Figure 1F), consistent with the previous demonstration that *rde-1* RNAi-defective mutations suppress the lethality of *adar-rrf-3* triple mutant (8). DRH-1 is thought to facilitate primary viral siRNA production from viral dsRNA by Dicer (18), and like the Argonaute RDE-1 is thought to be almost exclusively involved in RNAi of exogenous and viral dsRNA, whereas the other suppressors of the lethality of the *adar-eri* triple mutant DCR-1, RDE-4 and MUT-16 are required in both exogenous and endogenous RNAi pathways. The Argonaute NRDE-3 binds secondary siRNAs produced by RdRPs to induce transcriptional gene silencing. Thus, *adar-eri* triple mutant is sickly because of the hyper-induction of anti-viral RNA interference response that includes a nuclear RNAi component.

How is the RNAi machinery induced? Using RNAi of *drh-1/RIG-I* and *nrde-3/AGO*, the *adar-eri* triple mutant phenotype was suppressed and mRNAseq was used to quantify the expression of other RNAi factors. As a control, *adar-eri* triple mutant animals were exposed to a control RNAi vector that produces a ~200 bp vector dsRNA that does not match the *C. elegans* genome. Compared to wild type worms not exposed to this control dsRNA, most RNAi factors are induced two-fold in wild type worms exposed to control RNAi, possibly because of the presence of the 200 bp dsRNA (Supplemental Figure 1H). As a consequence, the induction of the RNAi machinery in the *adar-eri* triple mutant on RNAi control vector compared to wild type on RNAi control vector is less pronounced than the induction observed without exogenous RNAi. However, our data show that the sum of the induction of the RNAi machinery by the control RNAi vector and as a consequence of the *adar-eri* mutations is dependent on the Dicer complex gene *drh-1/RIG-I* and on the siRNA-binding protein *nrde-3/AGO* (Figure 1G, Supplemental Figures 1I and 1J), suggesting that the induction of the RNAi machinery requires accumulating siRNAs and that dsRNA is not sufficient to induce the RNAi machinery.

In summary, in the absence of ADAR editing, and with a hyperactive RNAi response because of the absence of the negative regulator of RNAi *eri-6/7*, the *adar-eri* triple mutant produces increased numbers of siRNAs resulting in severe phenotypes, that depend on the canonical RNAi pathway as well as the nuclear RNAi pathway. Both these pathways are induced and this induction depends on the viral RNA sensor DRH-1 and the nuclear RNAi factor NRDE-3. Whereas DRH-1/RIG-I was previously shown to act in exogenous RNAi and anti-viral RNAi, these data establish a role for DRH-1 in endogenous RNAi.

### Editing of palindromic repeat RNA affects gene expression

Since ADARs and RNAi act on dsRNA, we analyzed the genes that produce these siRNAs with respect to the double-stranded nature of their transcripts using published data of dsRNA purified using an anti-dsRNA antibody (19). One fifth of the siRNA-producing loci produce dsRNAs in wild type (Figure 2A). In the absence of ADAR activity, the dsRNA content of the cell is likely higher, since A->I edits destabilize dsRNA structures because A-U base pairs are replaced by I-U wobble base pairs.

**Figure 2.**
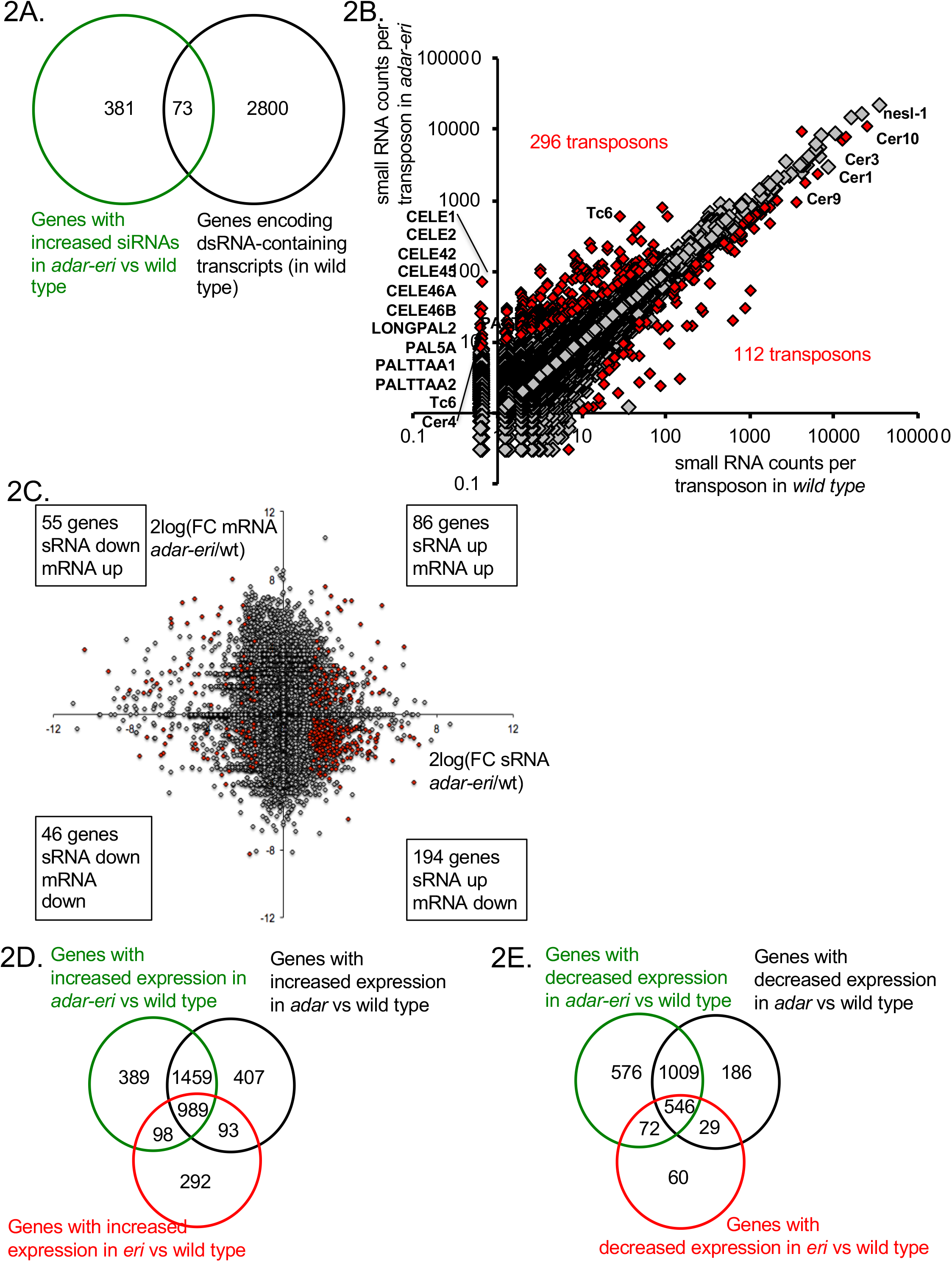
A. dsRNA-containing transcripts produce siRNAs in *adar-eri* triple mutants. B. siRNAs originate from transposons in *adar-eri* triple mutants. C. mRNA Expression changes in adar-eri/wt for each gene in the genome of versus changes in siRNA. P-value cut-off = 0.1. D. Overlap between genes silenced by the *eri, adar* and *adar-eri* genes. E. Overlap between genes silenced in the absence of the *eri, adar* and *adar-eri* genes

A source of dsRNA are palindromic repeat elements and inverted-repeat-containing transposons that have inserted into genes and are therefore expressed. We mapped the small RNA to ~11,000 transposons and ~65,000 inverted repeats. 296 transposons and 487 inverted repeats produce more siRNAs in the *adar-eri* triple mutant than wild type (Figure 2B and Supplemental Figure 2A). The transposons with increased siRNAs in the *adar-eri* triple mutant generally do not encode transposase proteins, have long inverted repeats and are predicted to form extensive hairpins (Supplemental Figure 2B).

How does this anti-viral RNAi response against palindromic dsRNA affect gene expression? siRNAs traditionally silence genes, which can be a consequence of either mRNA slicing by Argonautes and/or transcriptional silencing by nuclear RNAi. For the genes with a two-fold or more increase in siRNA expression in the *adar-eri* triple mutant, two thirds have significantly decreased expression in our total RNAseq analysis (Figure 2C). In an analysis of all gene expression changes (independent of editing or siRNA changes) we found that ADAR and ERI-6/7 both target many of same genes that are silenced in wild type (Figure 2D). This suggests that ERI-6/7 RNAi pathway target genes may have dsRNA structures that directs them to the RNAi pathway. In addition to the observed overlap between genes silenced by ADAR and ERI-6/7, there is also an overlap of genes with reduced expression in ADAR and ERI-6/7 versus wild type (Figure 2E, Supplemental Figures 2C and D and Supplemental Table 1).

In summary, palindromic repeat-containing genes produce siRNAs in the *adar-eri* mutant. In wild type, these palindromic repeat dsRNAs are edited (Supplemental Figures 2E-H). Editing could result in a destabilization of the hairpin; in the absence of editing, these hairpins are a substrate of Dicer, and produce siRNAs. Thus, ADAR editing, together with the *eri-6/7* RNAi pathway, prevents an RNAi response against self.

### ADARs and ERI-6/7 regulate LTR retrotransposons and endogenous viral elements

Among the mis-regulated transcripts in the *adar-eri* triple mutant are two transcripts corresponding to two orthologous genes of unknown function, C38D9.2 and F15D4.5; these transcripts are up to 60-fold induced in the *adar-eri* triple mutant versus wild type or *eri-6/7* (Figure 3A and C, Supplemental Table 2), with a slightly lower induction in the *adar* double mutant versus wild type, and no induction in the *eri-6/7* mutant. These genes (henceforth named Cer19-ORF2) are adjacent to annotated Cer19 LTR retrotransposon gag-pol genes and are likely part of these Cer19 retrotransposons: the genes are found in between the LTRs of Cer19 and encode proteins that have zf-CCHC gag-knuckle signatures typical of nucleocapsid proteins (Figure 3B, Supplemental Figure 3A). These Cer19-ORF2 copies encode expressed proteins (20).

**Figure 3.**
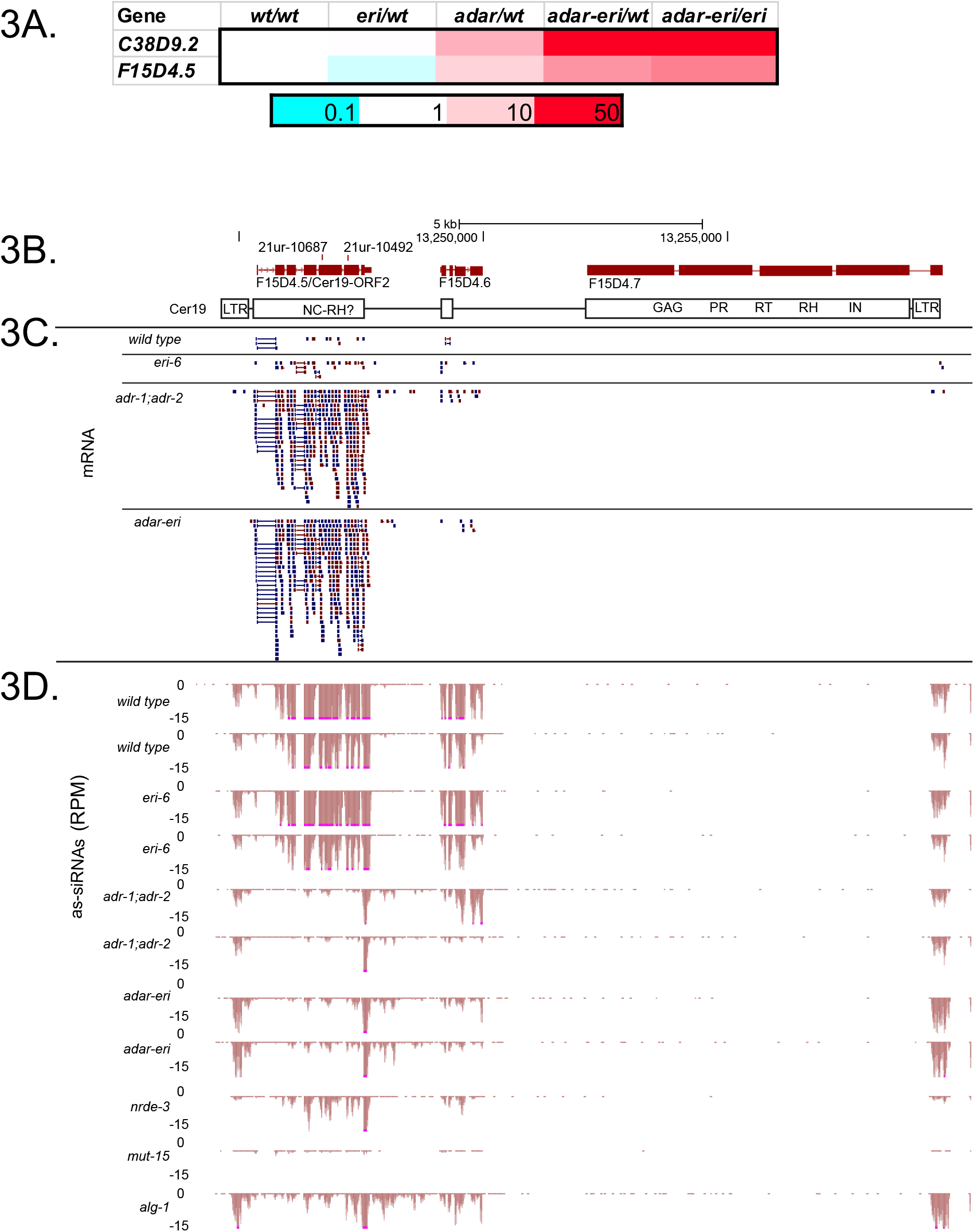
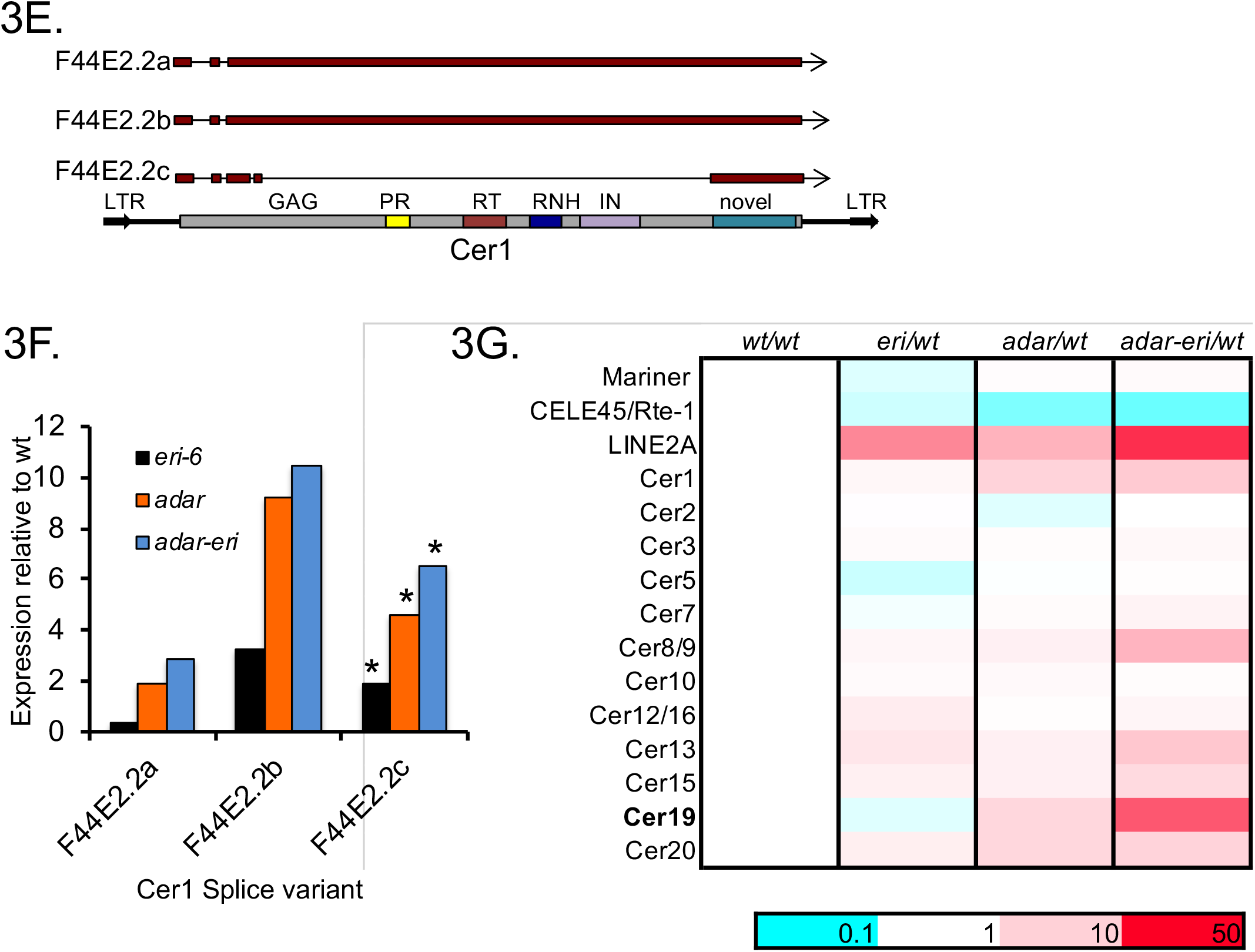
Cer19 retrotransposon sequences are upregulated in *adar-eri* triple mutants versus wild type. A. Two orthologous genes are highly upregulated in *adar-eri* triple mutants; indicated is the fold change versus wild type (P<0.01). B. The Cer19 retrotransposon genome is segmented. C. Total RNAseq shows an increase in Cer19 expression in *adar-eri* triple mutants. Shown are sequence reads. D. Anti-sense (as) siRNAs are depleted in *adar-eri* mutants. Duplicates are shown for wild type, *eri-6, adar* and *adar-eri*. Peaks are cut-off at 15 reads per million (indicated in pink). E. Cer1 transcripts and protein domains. Exons are indicated in red. F. Expression of Cer1 transcripts in *adar-eri* triple mutants relative to wild type (P<0.01). G. Expression of retrotransposons, and a DNA transposon (*mariner)* in *adar-eri* triple mutants relative to wild type. Cer1-20 are LTR retrotransposons.

Although *C. elegans* has relatively few full-length retrotransposons (14, 15), the genomes of related nematodes *C. brenneri* and *C. inopinata* contain an expansion of transposable elements (Supplemental Figure 3B) (21). These nematodes have retrotransposons homologous to Cer19 that contain at least two genes: ORF2, and a gag-pol gene (Supplemental Figure 3A). The closest *C. inopinata* Cer19 homolog encodes an envelope protein in addition to these two genes, suggesting that Cer19 is an endogenous retrovirus in *C. inopinata*.

For one *C. elegans* retrotransposon evidence of activity exists, although new insertions have never been observed the wild type strain that is widely used in *C. elegans* research: the Cer1 Ty3/gypsy element (Figure 3E) (22). VLPs of the Cer1 have been observed, and an insertion of Cer1 into a coding gene exists in some strains of *C. elegans* that is not present in wild type. Like Cer19, Cer1 RNA is expressed at increased levels (11-fold) in *adar-eri* triple mutants versus wild type (Figure 3F, Supplemental Figure 3C, Supplemental Table 2). Whereas many full length LTR retrotransposons are regulated by ADAR and ERI-6/7 (Figure 3G), other classes of transposons are not mis-regulated in *adar-eri* mutants, showing that LTR retrotransposons are differentially regulated from other types of transposons.

We previously identified ~100 genes that are silenced by the ERGO-1-ERI-6/7 RNAi pathway; these genes are likely recently integrated viruses because of their poor conservation between nematode species, repeated integrations in the worm genome and gene structure (7). Although the lack of conservation of these genes relegates them to molecular genetic nugatory, *C. elegans* produces high numbers of ERI-6/7-dependent siRNAs corresponding to these genes (Supplemental Table 3). These siRNAs may provide protection against future infections with closely related viruses. Several of the genes that the ERI-6/7 pathway silences encode viral envelope proteins homologous to the phlebovirus glycoprotein G2 that is found in Cer19 elements in other *Caenorhabdites* (Supplemental Figure 3D). Notably, *Caenorhabditis inopinata* has lost *eri-6/7* and *ergo-1* (21) and has, in addition to an expansion in retrotransposons, a large expansion of the number of viral envelope protein genes (Supplemental Table 4); the loss of the ERI-6/7 pathway could be the cause of this viral invasion or expansion of the *C. inopinata* genome. In addition, a gene encoding a protein that has homology to viral DNA polymerases present in high copy numbers in other *Caenorhabdites,* is also silenced by *eri-6/7* dependent siRNAs (7) (Supplemental Figure 3E). Thus, whereas the ADARs together with ERI-6/7 silence LTR retrotransposons, the ERI-6/7 pathway silences integrated viral genes that may be part of endogenous retroviruses.

### ADARs and the ERI-6/7 helicase are required for retrotransposon siRNAs

Since the ERI-6/7 helicase acts in RNAi targeting recently acquired genes, we hypothesized that Cer19 retrotransposon expression is regulated by siRNAs. Small RNA sequencing showed that in wild type, Cer19-ORF2 is ranked among the top thirty siRNA-producing genes in the genome (Supplemental Table 3), and siRNAs matching to Cer19-ORF2 are up to fourteen-fold depleted in the *adar-eri* triple mutant (Figure 3D, Figure 4A). siRNAs corresponding to the LTRs are not depleted (Figure 4B). The siRNAs are of 22 nucleotide length and have a 5’G typical of secondary *C. elegans* siRNAs produced by RNA-dependent RNA polymerases (Figure 4C). For the LTR retrotransposons Cer1 and Cer9 (Figures 4A, Supplemental Figure 3F) anti-sense siRNAs are partially depleted in the *adar-eri* triple mutant. This suggests that LTR retrotransposons are silenced by siRNAs which are dependent on the activity of ADARs and ERI-6/7. Retrotransposons are not obvious candidates of ADAR regulation, since ADARs lack inverted repeats that could produce dsRNA. Exogenous siRNA production is initiated at sites of cleavage in the mRNA that are subsequently uridylated before serving as templates for RdRP activity (23). In endogenous RNAi there is no dsRNA present that directs the slicing of the mRNA through primary siRNAs. It is not known which features flag an endogenous gene for siRNA production by RdRPs. In the *adar* double and *adar-eri* triple mutants, siRNAs are depleted for the 3’UTR and ORF; however, siRNAs downstream of the 3’UTR are increased suggesting that the RdRP that produces these types of antisense siRNAs may initiate at inappropriate sites downstream of the 3’UTR in these mutants. Using RNAfold to predict secondary structures in the 3’UTR and flanking sequences, we found that the polyadenylation signal sequence of Cer19-ORF2 is predicted to fold into a hairpin (ΔG=−4.7). A similar hairpin structure that includes the polyadenylation signal in retroviruses inhibits polyadenylation. Loss of destabilization of this hairpin as a result of absence of ADARs or the ERI-6/7 helicase could affect 3’end processing/cleavage, and thus the ability of the RdRP to initiate siRNA production.

**Figure 4.**
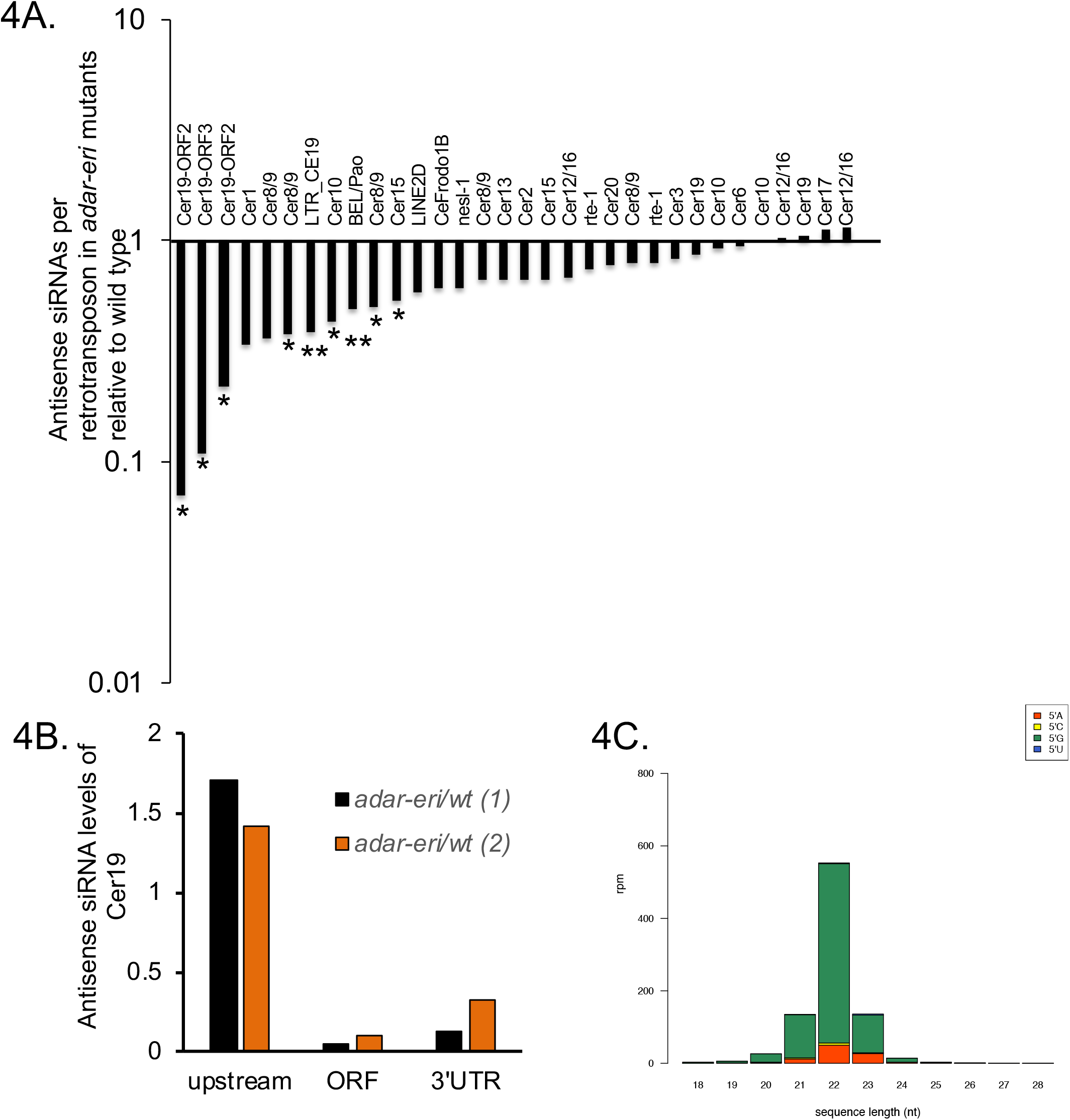
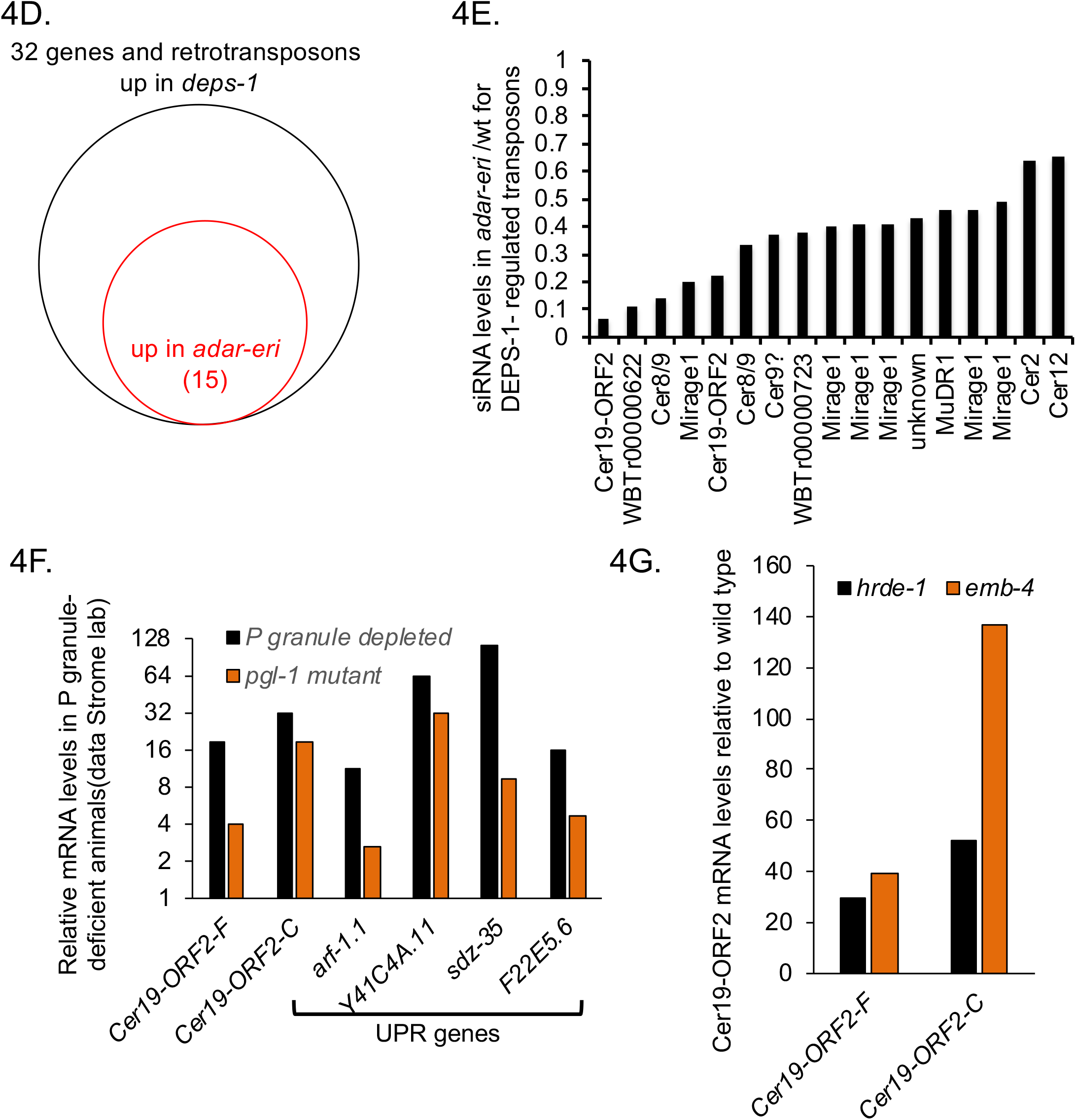
P granule components and RNAi factors silence retrotransposons. A. Relative levels of antisense siRNAs of full length retrotransposons in *adar-eri* triple mutants versus wild type. *: P<0.05, **: P<0.1. B. Antisense Cer19 siRNAs corresponding to ORF2 and the 3’UTR but not the 5’UTR and promoter are depleted in *C. elegans.* C. Antisense Cer19 siRNAs in wild type are predominantly 22G secondary siRNAs. D. Many DEPS-1-regulated genes are with ADAR-ERI-regulated retrotransposons E. Antisense siRNAs are decreased in *adar-eri* triple mutants versus wild type for DEPS-1-regulated retrotransposons. F. Animals lacking P granules have increased expression of Cer19 (data Strome lab). G. Cer19 is silenced by the nuclear Argonaute HRDE-1 and its interactor EMB-4 (Data from Akay *et al.)* (P<0.01).

DNA transposons are typically silenced by piRNAs that trigger siRNA generation. Generally, piRNAs are not depleted in the *adar-eri* triple mutant (Figure 1D). However, the Cer19 elements are predicted to be targeted by piRNAs (21ur-10687 and 21ur-10492) that are encoded on the opposite strand of one of the Cer19 elements (Figure 3B). These piRNAs are strongly depleted in *adar-eri* and *adar* mutants, suggesting that ADARs affect the biogenesis or stability of piRNAs, and thus the generation of siRNAs. However, the predicted piRNAs lack features typical of piRNAs: they do not have a 5’U, they are not dependent on the PIWI protein PRG-1, they are completely dependent on mutator proteins (suggesting that these small RNAs are siRNAs) (e.g. *mut-15* in Figure 3D), and in *prg-1* mutants, the target of these piRNAs, Cer19 RNA is two- to five-fold downregulated (piRTarBase (24)).

### Retrotransposon silencing requires nuclear RNAi factors and P granules

DNA transposons in *C. elegans* are silenced in the germ line through a process that involves scanning of mRNAs exiting the nucleus in perinuclear P granules to identify transcripts that need to be silenced (*i.e.* targets of piRNAs). A critical component of P granules is the DEPS-1 protein that is conserved in nematodes (25). Spike et al (2008) identified 32 genes that are up-regulated in the *deps-1* mutant. We found that at least twenty of these genes are retrotransposons and include the Cer19-ORF2 genes. Many of the DEPS-1-regulated retrotransposons are dramatically upregulated in the *adar-eri* triple mutant and depleted for siRNAs in the triple mutant (Figures 4D and 4E, Supplemental Figure 4A). DEPS-1 positively regulates the RNAi factor *rde-4/dsRBD*, and many DEPS-1- and ADAR-ERI-regulated retrotransposons are also regulated by the RNAi factors *mut-2/TENT* and *rde-4/dsRBD* (Supplemental Figure 4B). Similarly, in *pgl-1* mutants that are also defective in P granules (26), retrotransposons are desilenced (Figure 4F).

To determine whether the ADARs and ERI-6/7 affect gross P granule morphology, we performed immunohistochemistry for the P granule component PGL-1 in *adar-eri* triple mutant embryos. We did not observe any gross defects nor a reduction in PGL-1 expression (as has been observed for *deps-1* mutants) in *adar-eri* triple mutants, suggesting that the absence of ADARs and ERI-6/7 does not disrupt P-granule production or cell biology at these stages (Supplemental Figure 4C). Since DEPS-1 regulates positively *rde-4*, and loss of *rde-4* results in retrotransposon desilencing, loss of P granules may affect retrotransposon silencing through loss of *rde-4*.

Downstream of mRNA scanning and siRNA generation, RNAi target genes are typically silenced by nuclear Argonaute proteins, HRDE-1 in the germline or NRDE-3 in the soma. HRDE-1 and its interactor EMB-4/Aquarius (27) (28) silence DEPS-1-regulated retrotransposons (Figure 4G, Supplemental Figure 4D). Loss of somatic nuclear Argonaute NRDE-3 and the viral RNA sensor DRH-1/RIG-I does not disrupt retrotransposon silencing (Supplemental Figure 4E). Finally, loss of NRDE-3 and DRH-1/RIG-I does not suppress the *de*silencing of retrotransposons in the *adar-eri* triple mutant (Supplemental Figure 4E). This indicates that overproduction of palindromic siRNAs in the *adar-eri* mutant through DRH-1 does not trigger retrotransposon desilencing.

Taken together, whereas the upstream factors in LTR retrotransposon silencing differ from silencing of DNA transposons with the involvement of ADARs, ERI-6/7 and RDE-4 versus piRNAs, both retrotransposon and DNA transposon silencing depend on downstream nuclear RNAi factors and require P granules.

### The UPR is induced when retrotransposons are desilenced in the absence of ADAR editing and ERI-6/7 anti-viral RNAi

To identify additional pathways that regulate retrotransposons, we analyzed genes co-expressed with the Cer19-ORF2 genes using the SPELL engine, a query-driven search engine using all published *C. elegans* mRNA expression data (29). One third of genes most similar in expression profiles to Cer19 are other retrotransposons, indicative of a central regulatory mechanism of retrotransposon expression or silencing (Figure 5A). Among the non-retrotransposon genes co-expressed with retrotransposons, are the UPR genes *sdz-35* (a *C. elegans* paralog of KCTD10, substrate-specific adapter of a BCR (BTB-CUL3-RBX1) E3 ubiquitin-protein ligase complex) and *rrf-2* (an RdRP), both of which are highly induced in conditions of protein misfolding induced by the drug tunicamycin (30).

**Figure 5.**
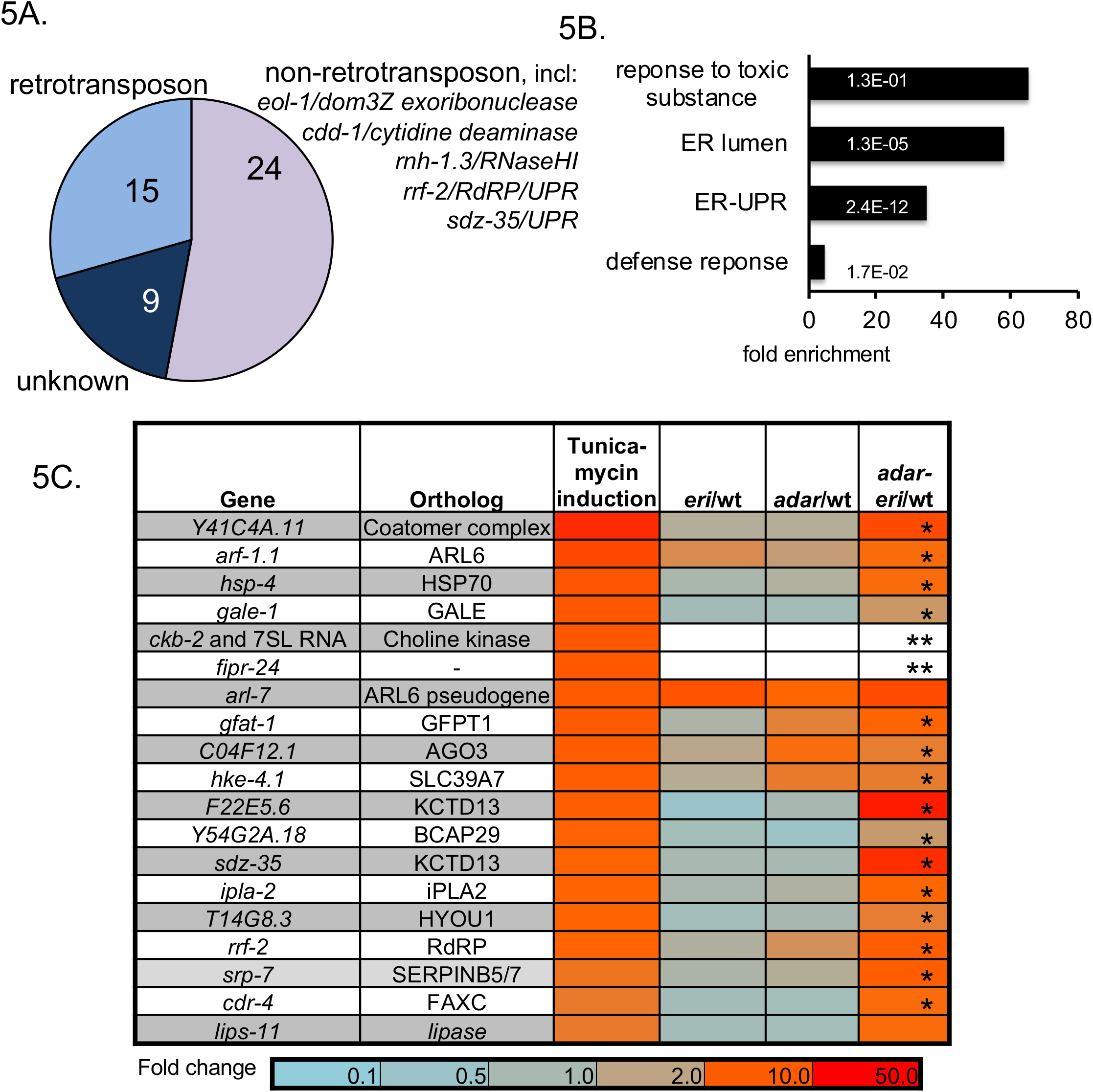
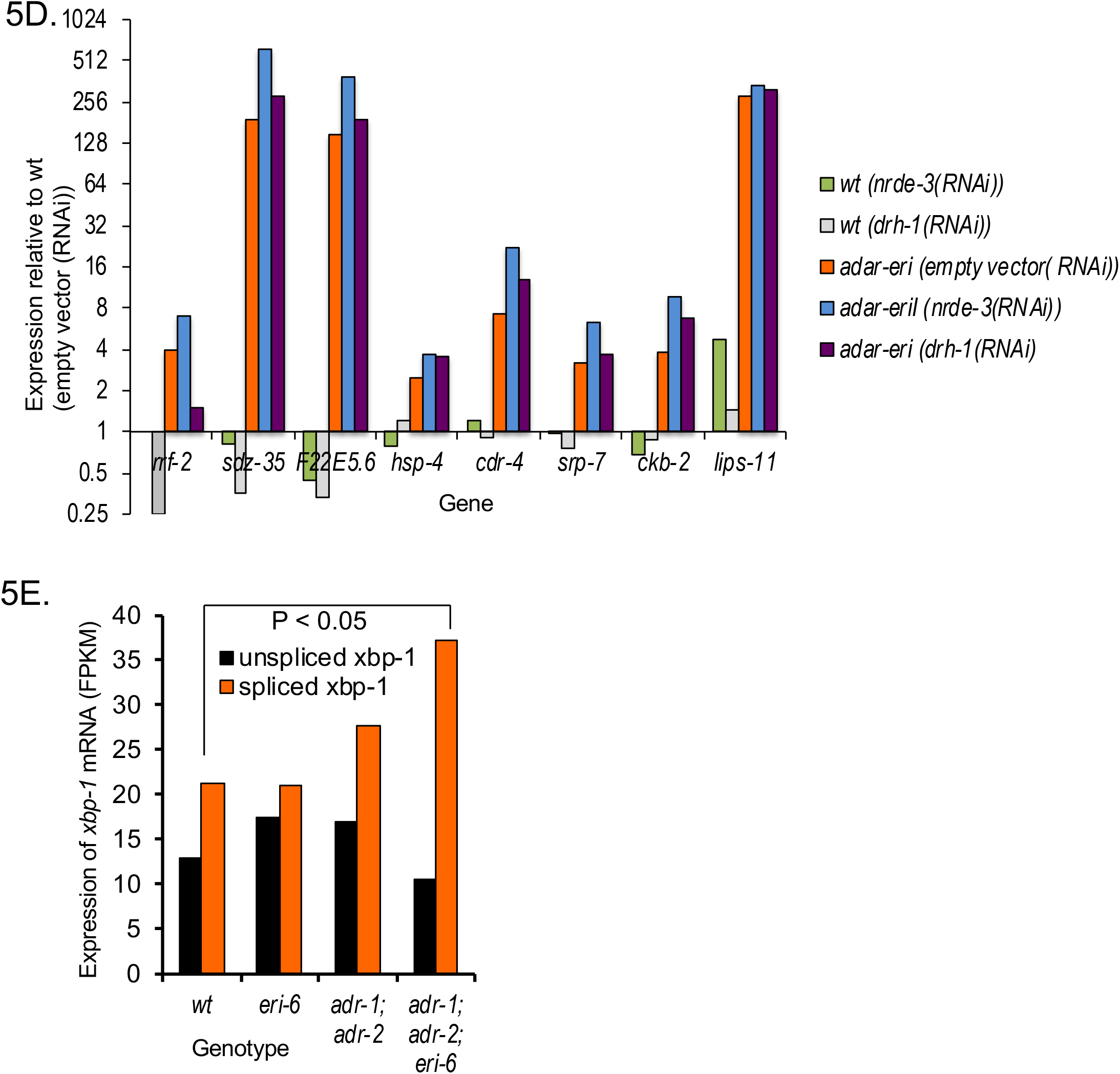
A. Coexpression analysis of Cer19. Analysis of the 50 most similarly expressed genes based on cumulative expression profiling (SPELL (Hibbs *et al.* 2007)) B. GO-term enrichment analysis of genes only upregulated *adar-eri* triple mutants (168 genes). C. Relative expression levels of UPR genes in *adar* and *eri* mutants (RNAseq) of the 19 most-induced genes when wild type *C. elegans* is treated with the UPR-inducing drug tunicamycin (data Kaufman lab). (* =significant P<0.01, ** not expressed in wild type but highly expressed in *adar-eri* triple mutants). D. and E. The induction of the UPR in *adar-eri* mutants is independent of the RIG-I ortholog DRH-1 (D) and the Argonaute NRDE-3 (E). The induction of the RdRP gene *rrf-2* is dependent on DRH-1. F. Expression of non-IRE-1-spliced *xbp-1* mRNA and IRE-1-spliced *xbp-1* mRNA in *adar* and *eri* mutants (RNAseq).

The *adar-eri* triple mutant shows a dramatic induction of genes with GO-terms related to the Unfolded Protein Response (UPR) relative to wild type (Figure 5B). Of the 20 *C. elegans* genes most induced by the ER stress-inducing drug tunicamycin (30), all are similarly induced in *adar-eri* triple mutants, but not in *adar* and *eri* single mutants (Figure 5C). To determine whether the induction of the UPR is a consequence of the hyperactive RNAi response that produces new siRNA from palindromes, we analyzed mRNAseq data of *adar-eri* triple mutants in which the RNAi genes *drh-1* and *nrde-3* that mediate the hyperactive RNAi response are depleted. These data show that these RNAi factors are not required for the induction of the UPR and also not for the desilencing of retrotransposons, showing that the developmental and fertility defects of the *adar-eri* triple mutant are independent of retrotransposon desilencing (Figure 5D).

The UPR consists of three branches, namely IRE1, PERK, and ATF6. A signature of the induction of the IRE1 branch is the IRE-1-mediated cleavage of the mRNA of the transcription factor XBP-1, excising an ‘intron’ resulting in a functional XBP-1 mRNA. An analysis of our RNAseq data, and confirmation using qRT-PCR, showed increased IRE-1-mediated excision of the *xbp-1* ‘intron’ in the *adar-eri* triple mutant (Figure 5E and Supplemental Figure 5A), indicating a bona fide UPR response involving the IRE1 branch.

The coexpression of retrotransposons and UPR genes suggests a link between the induction of the UPR and the over-expression of retrotransposons observed in the *adar-eri* triple mutant. Like retrotransposons, UPR genes are upregulated in animals deficient for P granules (Figure 4F). Viruses use the ER for maturation and assembly of virus particles (16). Similarly, the yeast retrotransposon Ty1 RNA is translated in association with the SRP to target the nascent peptide to the ER. The Ty1 gag protein is subsequently translocated into the ER, a translocation that is necessary for Virus-Like Particle (VLP) assembly (17). Given the high overexpression of retrotransposon mRNAs and correlated UPR induction, it is possible that the overexpression of retrotransposons results in the accumulation of gag protein or Virus-Like Particles (VLPs) and that the induction of the UPR is caused by viral or retrotransposon protein-induced ER stress. To test this hypothesis we depleted the *adar-eri* triple mutant of Cer19 ORF2 RNA (one of the most highly overexpressed genes), and assayed for UPR induction (Supplemental Figure 5B). We observed a reduction in UPR gene expression in the *adar-eri* triple mutant depleted for Cer19-ORF2.

To study the interaction between RNAi factors, the ADARs and UPR genes, we assayed animals with mutations or RNAi depletions for multiple of these genes for potential interactions. We found interactions with mutations in the ER chaperone gene *hsp-4/BiP*. Either *adr-1* and *eri-6/7* mutations enhance the *hsp-4* mutant phenotype: *hsp-4 adr-1* double mutants are sterile at 15°C, *hsp-4; eri-6* double mutants are larval lethal at 15°C, whereas the *hsp-4* single mutant is viable at 15°C (Supplemental Figure 5C).

The drug tunicamycin inhibits N-glycosylation and thus induces the UPR. In *adar-eri* mutant animals, the UPR is constitutively induced, and thus an additional load of misfolded proteins may cause increased lethality. We tested the *adar* and *eri* mutants for tunicamycin sensitivity at 25°C. The *adar-eri*, *eri* and *adar* mutants were all more sensitive to 10 mg/ml tunicamycin than wild type (Supplemental Figure 5D), indicating the presence of an increased load of misfolded proteins in *adar* and *eri* mutants.

## *hsf-1/HSF1* regulates UPR genes and retrotransposons

Viral infection and retrotransposon activation induce a common transcriptional response: 13% of the genes induced specifically in the *adar-eri* mjutant (but not in the *adar* or *eri* mutants) are among the 320 genes induced upon infection with Orsay virus, a positive strand RNA virus that infects the intestine of *C. elegans* (Figure 6A, Supplemental Table 5). In addition, more than a third of the genes specifically upregulated in the *adar-eri* mutant are silenced by *pals-*22, a negative regulator of the Intracellular Pathogen Response (IPR) to the microsporidium *N. parisii*, indicating an overlap between the responses to retrotransposons and to intracellular pathogens (Figure 6A).

**Figure 6.**
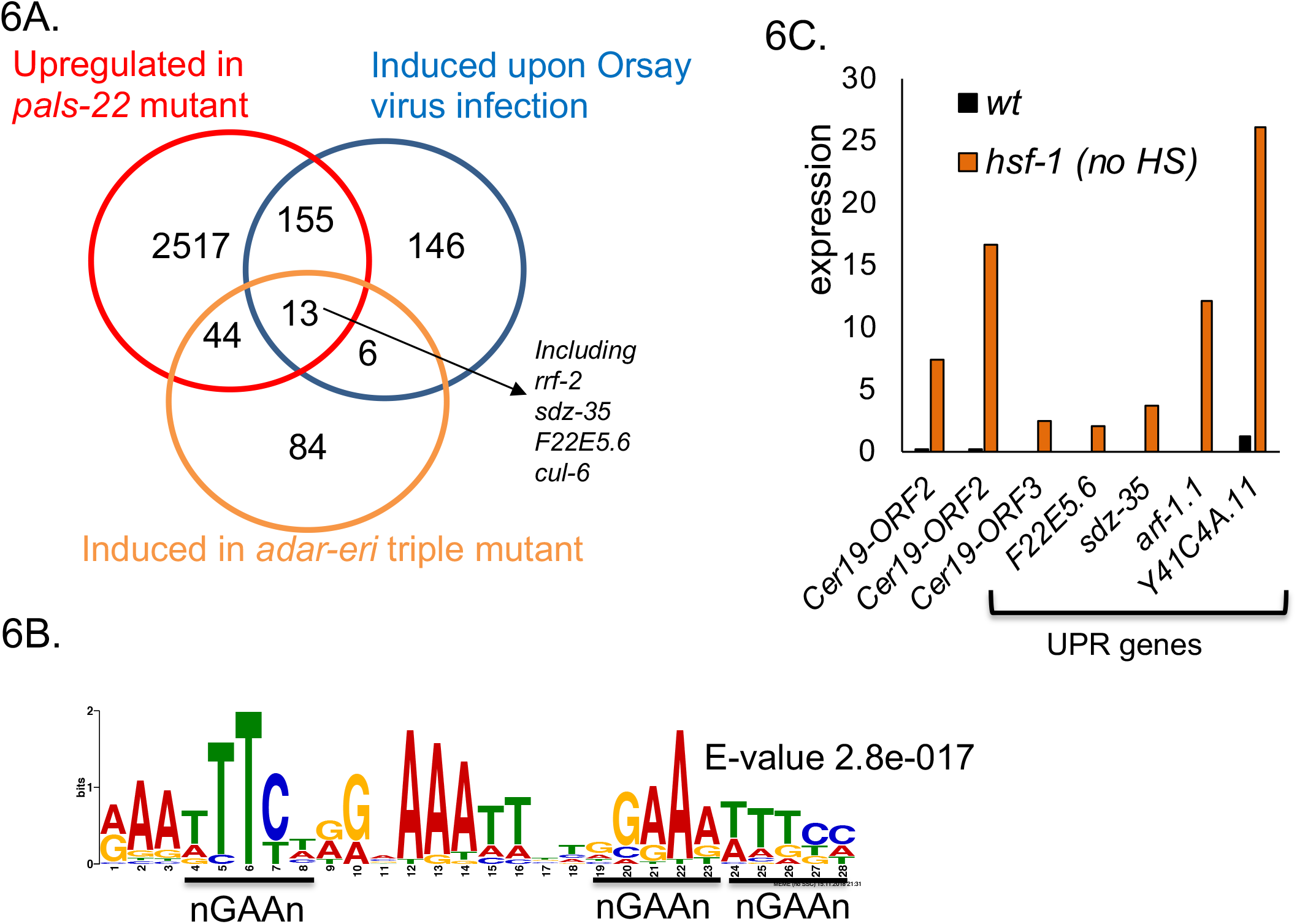
A. Genes induced in *adar-eri* mutants overlap with genes induced upon Orsay virus infection and genes upregulated in a *pals-22* mutant. B. A motif similar to the HSF-1 binding element (HSE) was identified in the 2 kb promoter regions of retrotransposons Cer19-ORF2 (2 copies of the motif) and UPR genes F22E5.6 (5 copies), *rrf-2*-(2 copies) and *sdz-35* (5 copies). C. Expression of retrotransposons and UPR genes in *hsf-1 (ok600)* mutant animals (without heat shock exposure) (mined data Morimoto lab).

To identify the pathways that regulate retrotransposons and UPR factors, we searched for and identified shared motifs in the promoter regions of these genes. A site similar to the HSF-1/heat shock factor binding site (HSE) was identified in the promoter regions of Cer19 ORF2 genes, and of several UPR genes (Figure 6B). Indeed, Cer19 and these UPR genes are developmentally regulated by HSF-1 (31, 32), with HSF-1 under physiological conditions repressing the expression of these genes (Figure 6C). The UPR genes *sdz-35* and *F22E5.6* are associated with HSF-1 and up-regulated in *hsf-1(ok600)*. The Cer19 genes and also many ERGO-1-ERI-6/7 targets are expressed exclusively at higher temperatures (at 25°C): of the 183 genes identified that only are expressed at 25°C and not at 15°C or 20°C (33), 33 genes are ERGO-1-ERI-6/7 targets (Supplemental Figure 5E). This increase in expression at 25°C is accompanied by a reduced expression of critical silencing factors *eri-6/7*, *emb-4* and *drh-3* (33). Cer19 retrotransposons are also expressed in response to heat shock (34). Heat shock leads to dsRNA accumulation in HSF-1 granules (35). In an RNAi screen for transcription factors that interact with the *adar-eri* triple mutant, we found that loss of *hsf-1* in the *adar-eri* triple mutant results in larval lethality (Supplemental Figure 5C), indicating that the loss of multiple retrotransposon silencing factors is lethal.

## DISCUSSION

ADARs in mammals prevent a RIG-I-mediated interferon response against endogenously encoded dsRNA, *e.g.* dsRNA produced from close insertions of two Alu elements into transcribed regions. Similarly, in *C. elegans,* ADARs, together with negative regulators of the RNAi pathway, prevent an DRH-1/RIG-I-dependent response triggered against genomically-encoded dsRNA-containing mRNAs, in this case an RNAi response. Many of these dsRNAs correspond to palindromic repeat elements that have inserted into genes. This anti-viral RNAi response against endogenous dsRNA has profound effects on the transcriptome mediated by the nuclear RNAi factor NRDE-3. (8). The Argonaute NRDE-3 acts in the ERGO-1 and ERI-6/7 endogenous RNAi pathway to silence viral genes, but also acts in the exogenous RNAi pathway to silence genes targeted by exogenous dsRNA. NRDE-3 shuttles siRNAs from the cytoplasm into the nucleus to direct histone modifications to the corresponding genes and trigger heritable gene silencing. In the ERI-6/7 mutant, the ERI-6/7-dependent siRNAs are missing, and NRDE-3 is cytoplasmic. Since loss of NRDE-3 suppresses the lethality of the ADAR-ERI triple mutant, it is likely that NRDE-3 binds the novel palindromic siRNAs (that are similar to exogenous siRNAs) and shuttles them into the nucleus to induce histone H3 modifications, ultimately causing lethality by inappropriate gene silencing.

The genomically-encoded dsRNAs trigger an increase in the expression of RNAi machinery factors (including *mut-16*, *rde-1*/AGO, *dcr-1/DICER)* in both the *adar* double mutants and in the *adar-eri* triple mutants. Both these mutants produce increased numbers of siRNAs. The RNAi-factor induction is dependent on DRH-1/RIG-I and NRDE-3/AGO, suggesting that is the overexpression of siRNAs that triggers this transcriptional response. One model to explain how the increased number of siRNAs is sensed in the cell is that endogenous siRNAs are out-competed by the new palindromic siRNAs for binding by NRDE-3 (or other Argonautes) thereby releasing silencing of an unknown regulatory factor that is usually silenced by endogenous siRNAs. Alternatively, an Argonaute that specifically binds exogenous (and palindromic) siRNAs and not endogenous siRNAs, like RDE-1, promotes RNAi machinery expression when in complex with siRNAs. The *eri-6/7*/MOV10 RNA helicase single mutant does not accumulate siRNAs derived from palindromic repeats, and therefore exogenous RNAi pathway genes are not induced. This enhanced RNAi (*eri*) mutant does have an enhanced response to many dsRNAs introduced by feeding *E. coli* expressing dsRNAs (7, 36). It is possible that when dsRNA from *E. coli* is introduced into the *eri-6/7* mutant, RNAi pathway genes are induced contributing to the enhanced RNAi phenotype, similar to the induction of the RNAi machinery in *adar* mutants by palindromic dsRNAs/siRNAs. In fact, the enhanced RNAi phenotype of an *eri-6/7* single mutant is suppressed by inactivation of many of the same RNAi components that suppress the inviability of the *adar-eri* triple mutant (7, 36).

While the ADARs and the ERI-6/7 RNAi pathway prevent silencing of palindromic repeats that can form double stranded RNAs, the ADARs and ERI-6/7 are required to silence retrotransposons and integrated viral genes. ERI-6/7 targets include viral envelope proteins and viral DNA polymerases that vary dramatically in copy number in different *Caenorhabdites,* with the *C. inopinata* genome haboring high copy numbers of retrotransposon genes and retroviral envelope genes. The lack of ERGO-1 and ERI-6/7 in *C. inopinata* supports a role for this pathway in silencing of viruses and retrotransposons. Loss of both ADARs and ERI-6/7 results in a strong upregulation of LTR retrotransposon expression, accompanied by a loss of siRNAs, particularly in the 3’UTR. Retrotransposons and ERI-6/7 target genes are among the genes that produce the most siRNAs in wild type worms. Why do ADAR-ERI mutants produce fewer retrotransposon siRNAs? Mov10 in mammals associates with TUTases that uridylate LINE-1 retrotransposons; uridylation prevents LINE-1 reverse transcription and destabilizes the RNA. Mov10 binds to 3’UTRs to displace 3’UTR proteins and resolve secondary structures. Possibly, loss of ERI-6/7/Mov10 results in an inability to resolve 3’UTR secondary structures that are also stabilized by the loss of ADARs. The presence of these structures may prevent siRNA generation because of an inability of an RdRP to target the 3’UTR, either because of lack of uridylation or presence of secondary structures. Alternatively, ADARs could affect dsRNA intermediates formed by small RNA primers base-paired with retrotransposon RNA that are generated during reverse transcription of LTR retrotransposons. Finally, ADARs could affect siRNA biogenesis independent of editing.

Downstream of siRNA biogenesis, retrotransposon silencing is dependent on P granule components and nuclear RNAi factors in the germ line, similar to silencing of DNA transposons downstream of piRNAs. We found that additional pathways affect retrotransposon expression: retrotransposons are de-silenced at elevated temperatures and require the HSF-1 heat shock factor for silencing, with potentially direct binding of HSF-1 to sites in the promoter regions of retrotransposons.

Desilencing of retrotransposons is accompanied by an upregulation of the unfolded protein response, in ADAR-ERI mutants but also in animals lacking P granules and in HSF-1 mutant animals. An study of the *adr-1;adr-2;rrf-3* mutant from the Bass lab(8) used poly(A)^+^ RNA-seq in embryos in their analyses; Cer19 and UPR genes are significantly up-regulated in these conditions, albeit not as dramatically. This difference is likely due to the analysis of a different in developmental stage of the animals (embryos versus adults) and the methodology (poly(A)^+^ versus total RNAseq).

In an analysis of genes coexpressed with Cer19 we found several UPR genes among genes co-expressed along with retrotransposons. The unfolded protein response is likely to be due to the increase in expression of viral proteins and/or virus like particle assembly at the ER. The genes that are induced in response to infection with Orsay virus, a non-enveloped positive strand virus, overlap with genes upregulated in *adar-eri* triple mutants, as well as with the genes that are co-expressed with LTR retrotransposons. All three conditions show an upregulation of the RdRP gene *rrf-2*, two genes encoding KCTD10 paralogs that are substrate-specific adapters of a BCR (BTB-CUL3-RBX1) E3 ubiquitin-protein ligase complex, the cullin gene *cul-6*, also a component of a E3 ubiquitin-protein ligase complex, and a gene encoding a protein with homology to PALS-26. *cul-6* acts in a stress response pathway to the intracellular pathogen *N. parisii*, called the intracellular pathogen response (IPR) that is induced upon proteotoxic stress caused by intracellular infection and or by prolonged heat stress, and is negatively regulated by *pals-22* (37). *pals-22* mutants also show upregulation of *rrf-2,* of the two genes encoding KCTD10 paralogs and of Cer19 (38) and many other genes specifically up-regulated in *adar-eri* mutants or coexpressed with retrotransposons. The identification of *cul-6/cullin* and substrate-specific adaptors of a E3 ubiquitin ligase complex suggests a role for ubiquitylation in the response to LTR retrotransposons in *C. elegans*. In mouse, LINE-1 retrotransposon mobilization is restricted by ubiquitylation of a retrotransposon-encoded protein by an E3 ubiquitin ligase, resulting in protein degradation in mouse embryonic stem cells (39)

The LINE2A retrotransposon is another retrotransposon that is silenced by the ADARs and ERI-6/7 (Figure 2G). QTL mapping of activity of the LINE2A retrotransposon in natural isolates of *C. elegans* has identified a number of natural gene variants that may be responsible for the loss of LINE2A silencing, including variants of the genes *adr-1/ADAR, rrf-2/RdRP, rnh-1.3/RNaseH1*, and *hsf-1/HSF1* (40). This finding supports the role of these genes in retrotransposon silencing.

*C. elegans* uses multiple mechanisms to silence repeats and transposable elements, many of which are conserved in human. Palindromic repeats are edited by ADARs, preventing an anti-viral RNAi response. DNA transposons are silenced by piRNAs. Integrated viral genes are targeted by the ERI-6/7/Mov10 endogenous RNAi pathway. LTR retrotransposons are silenced by ADARs together with the ERI-6/7/Mov10 pathway. The presence of many copies of viral envelope genes in *Caenorhabdites*, including within LTR retrotransposons, suggests that some of these may represent endogenous retroviruses. How these elements are targeted for silencing remains unknown. In addition to the diverse silencing mechanisms, there are similar responses in *C. elegans* to infection with a non-enveloped RNA virus, infection with an intracellular pathogen and to de-silencing of retrotransposons that are likely the result of proteotoxicity.

## MATERIALS AND METHODS

### *C. elegans* strains and culture

The following strains were used:

BB4 (*adr-1(gv6) adr-2(gv42)*), BB21 ((*adr-1(tm668) adr-2(ok735)*), GR1744 (*adr-1(gv6) adr-2(gv42) eri-6(mg379)*), SEF298 (*ergo-1 (tm1860) adr-1(tm668) adr-2(ok735)*).

Brood size assays were performed at 15, 20 and 25°C. For tunicamycin assays, embryos were dropped on plates containing 10 mg/ml tunicamycin and followed for arrest and lethality at 25°C. All other assays were performed at 20°C.

For mRNAseq, GR1744, N2, *eri-6*, and BB21 were exposed to feeding RNAi of *nrde-3, drh-1* or control vector dsRNA. For all mRNA, total RNA and small RNA sequencing experiments, worms were grown to 70 hrs after dropping L1-arrested worms onto *E. coli* OP50 at 20°C, the young adult stage that produces embryos.

### Immunostaining

PGL-1 immunostaining was done as described previously (41).

### qRT-PCR analysis UPR genes

cDNA was made using Retroscript kit with the following primers: (xbp-1: 1105 (ccgatccacctccatcaac) and 1106 (accgtctgctccttcctcaatg), 1107 (tgcctttgaatcagcagtgg) and 1108 (accgtctgctccttcctcaatg)); *act-1* control: 1113 (cttgggtatggagtccgcc) and 1114 (ttagaagcacttgcggtgaac); *Y45F10D.4* control 653 (gtcgcttcaaatcagttcagc) and 654 (gttcttgtcaagtgatccgaca); *srp-7* (aatgtctccagtacttcggttaatg and aattccgagcgattgaagag); *Y41C4A.11* (gccatggattttgactgctt and cgtggatttttcggagacc).

### RNAi suppression screen

SEF298 and GR1744 animals were fed on a library of *E. coli* producing dsRNA corresponding to genes encoding RNAi factors. Feeding (in duplicate) was started at the L1 stage; phenotypes were scored in adults of the next (F1) generation.

Suppression was scored as ‘strong’ when no rupture was observed, and as ‘moderate’ when fewer that 10% of progeny ruptured. Retests were done twice in duplicate in parallel with controls N2 (wt), eri-6, BB4, BB21. RNAi clones were sequenced to confirm gene identity.

### Small RNA differential expression analysis

Small RNA libraries were prepared as described before. Small RNA was mapped to the *C. elegans* genome (WBcel235) using Bowtie. Based on overlap between map positions of the small RNA, and the coordinates of microRNA and piRNA genes, microRNAs and piRNAs were identified. The remaining small RNAs are endogenous siRNAs. The siRNAs were mapped to genes (WS247 and WS260). The gene identities were used to further classify siRNAs into specific endogenous RNAi pathways mediated by particular Argonaute proteins. Small RNA reads were also mapped to genes (over 46,000), transposons (over 11,000), inverted repeats (over 65,000) using annotation files downloaded from Wormbase (10/2017) using bwa and featureCounts and analyzed using Deseq2.

### Total and mRNA differential expression analysis

rRNA was removed from total RNA using Ribo-zero (Epicentre/Illumina). RNAseq libraries were made using NEBNEXT Ultra. Single-end sequencing runs of 50 nts were done. Reads were first mapped to rRNA and tRNA using Bowtie2. Reads not mapping to rRNA and tRNA were then mapped to WBCel235 genome/WS247 annotation using Tophat2 with minimum intron size=20 (−i), maximum intron size 50,000 (−I) and allowing for 1 mismatch (−N 1). Both guided and *de novo* transcriptome assembly were performed, to identify new genes and transcripts. Differential expression analysis was done using Cufflinks, Cuffmerge and Cuffdiff with multi-read correct. To map to transposons and inverted repeats, featureCounts was used counting all alignments, Deseq2 was used to assess differential expression. A P value <0.1 was used as a cutoff for differential expression.

mRNAseq was done using NEBNext Poly(A) Magnetic Isolation Module with NEBNext Ultra Directional RNA library kit.

GO-term enrichment was analyzed using GOrilla.

## ACKNOWLEDGEMENTS

We would like to thank Peter Breen and Alexia Hwang for technical assistance. We also would like to thank members of the Ruvkun lab and Noémie Scheidel for suggestions.

