## Supplemental Figures and Tables for "*Caenorhabditis elegans* ADAR editing and the ERI-6/7/MOV10 RNAi pathway silence endogenous viral elements and LTR retrotransposons"

### Supplemental Figure 1

1A.

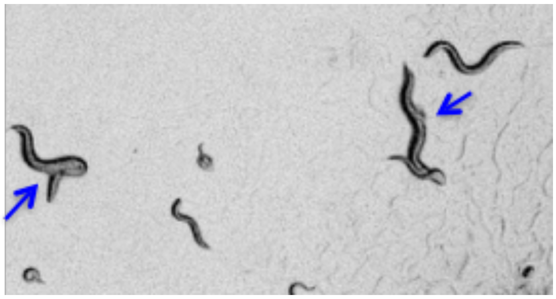

1B.

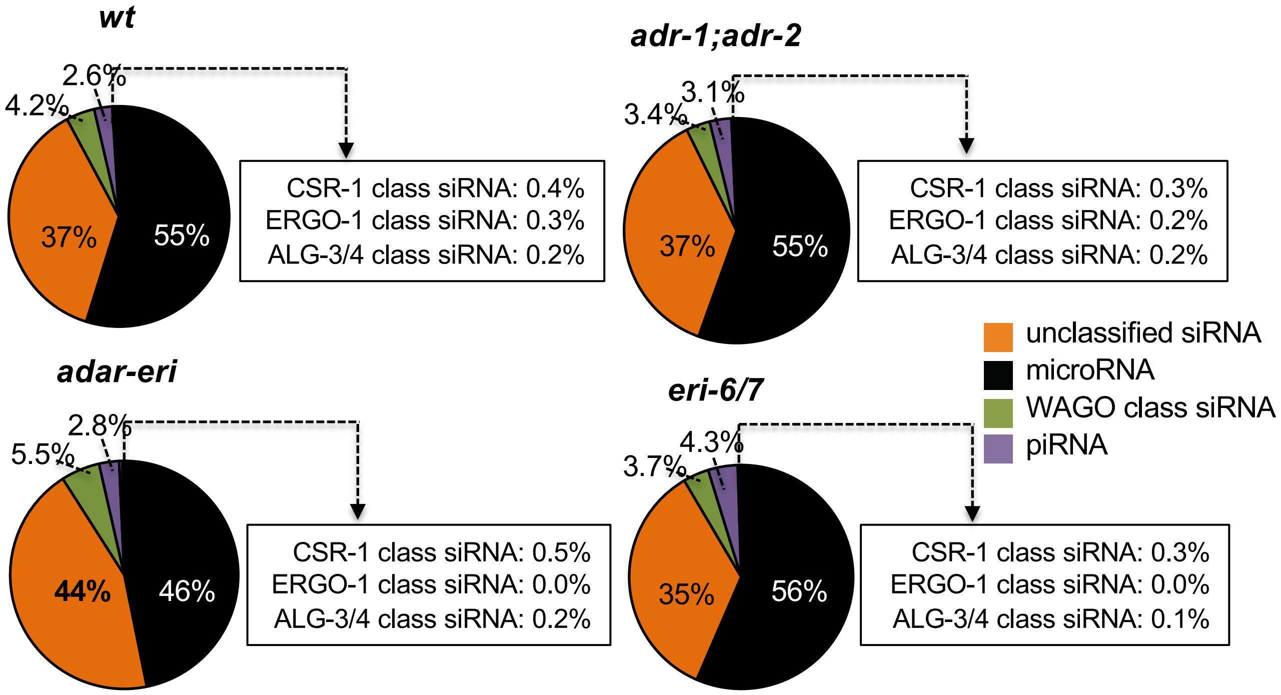

Supplemental Figure 1

1C.

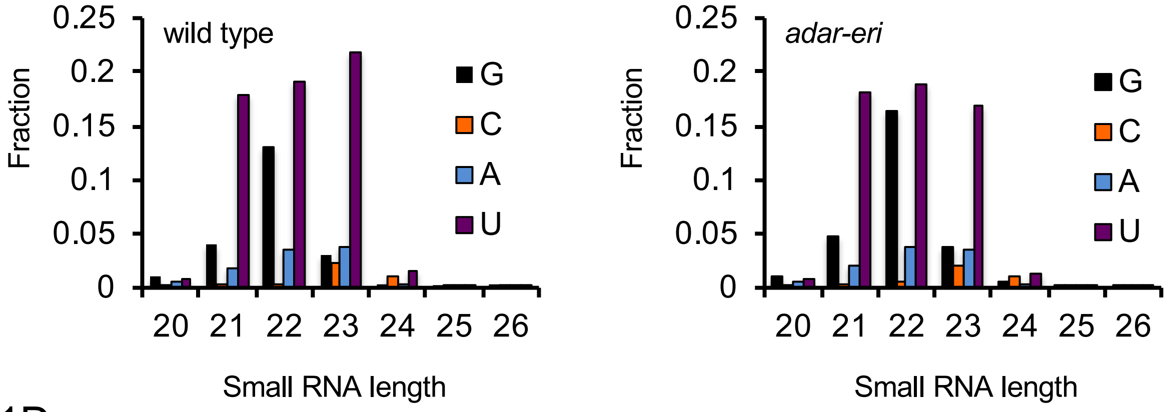

1D.

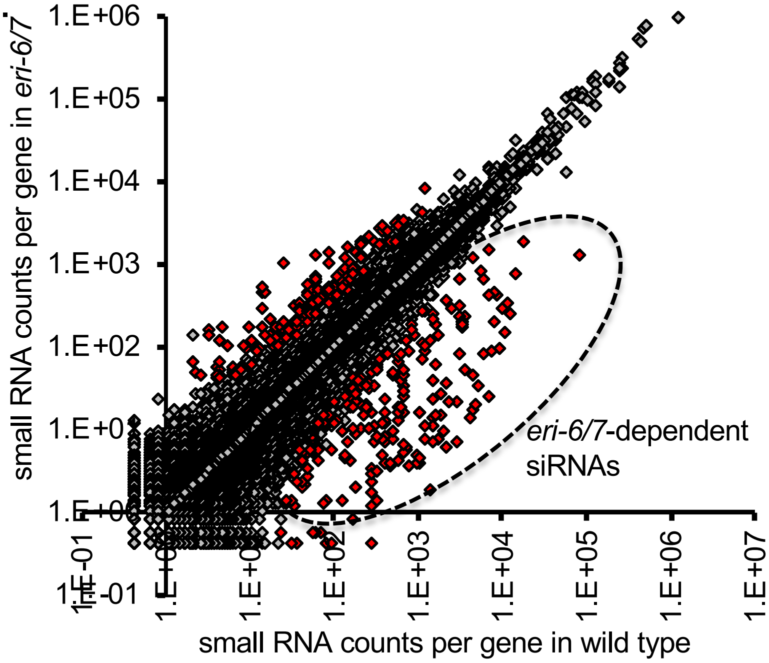

1E.

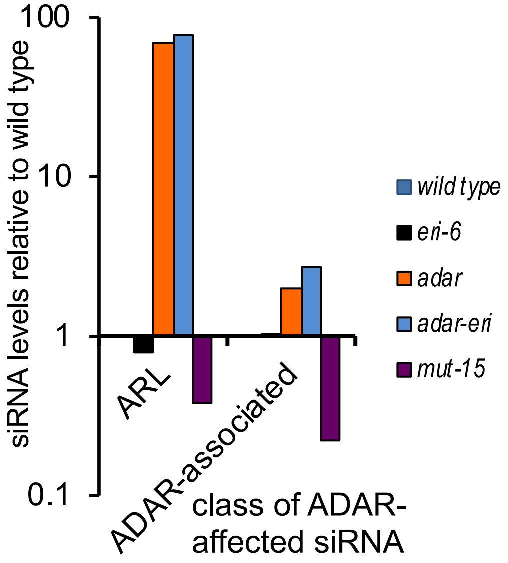

1F.

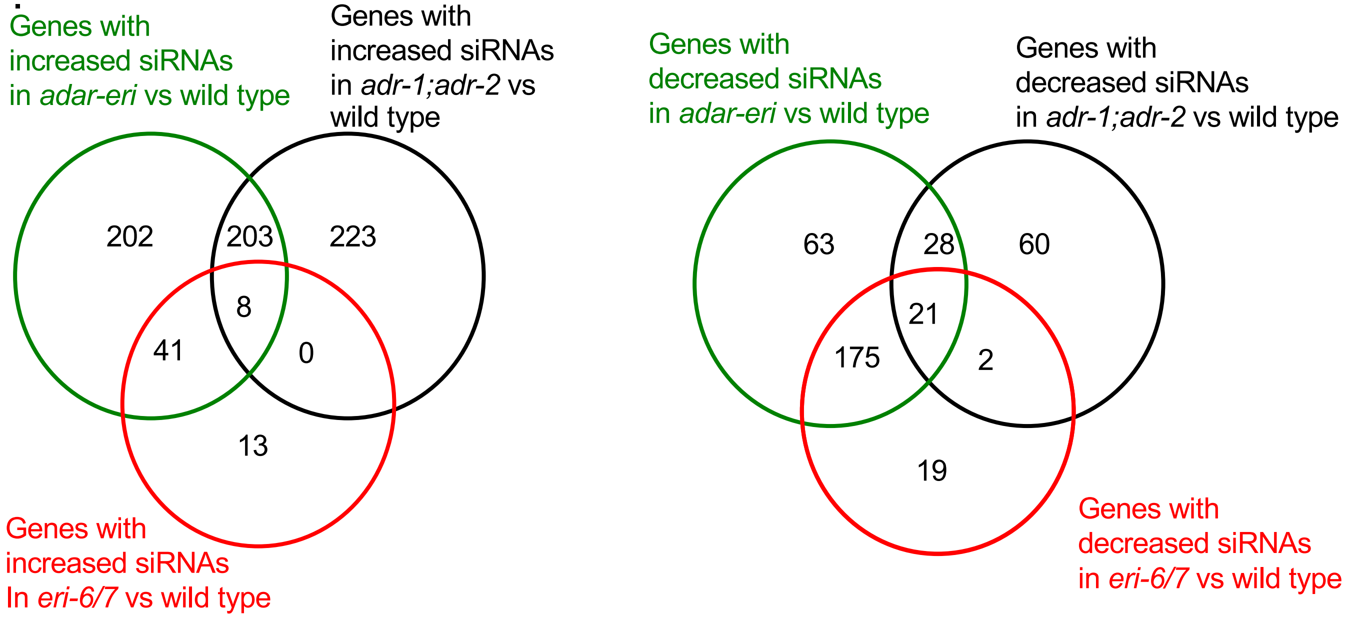

Supplemental Figure 1.

1G.

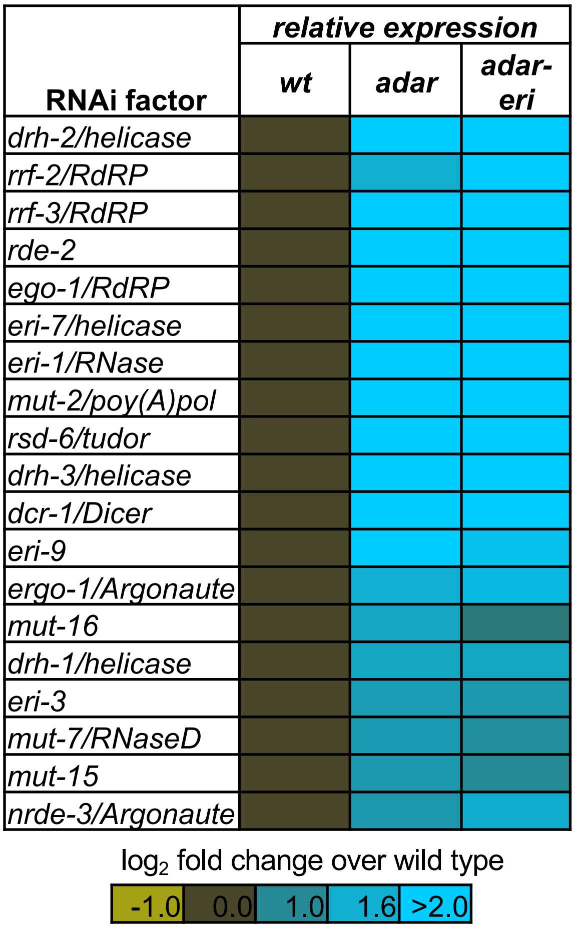

1H.

| RNAi factor | Expression (wild type on control RNAi/wild type without RNAi) |
|---|---|
| *eri-7/helicase* | 2.2 |
| *eri-1/RNase* | 1.8 |
| *dcr-1/Dicer* | 1.7 |
| *eri-9* | 1.8 |
| *ergo-1/Argonaute* | 1.8 |
| *eri-3* | 1.5 |
| *hrde-1/Argonaute* | 1.9 |
| *rde-1* | 1.8 |
| *nrde-3/Argonaute* | 1.6 |
| *nrde-2* | 1.8 |
| *rde-4/dsRBD* | 1.6 |
| *rde-10* | 1.5 |
| *nrde-1* | 1.9 |
| *wago-1/Argonaute* | 1.9 |

1I.

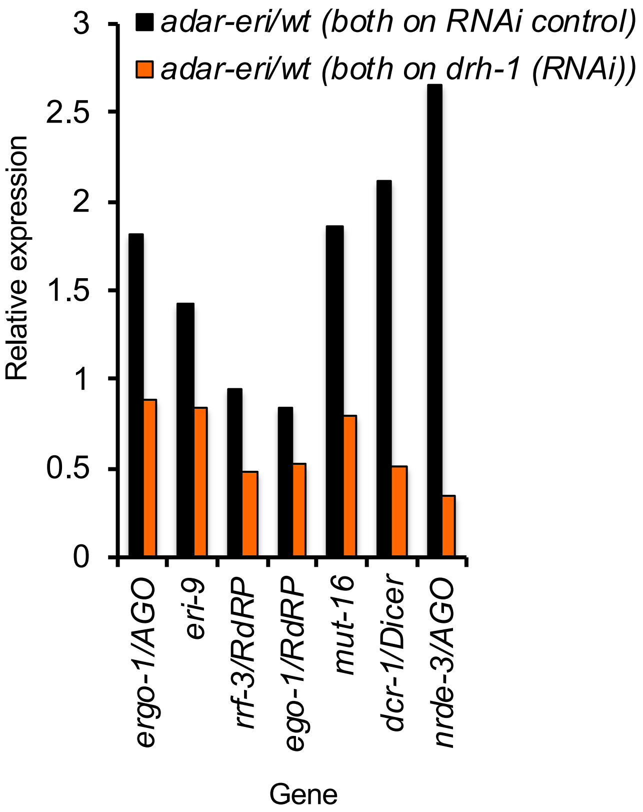

1J.

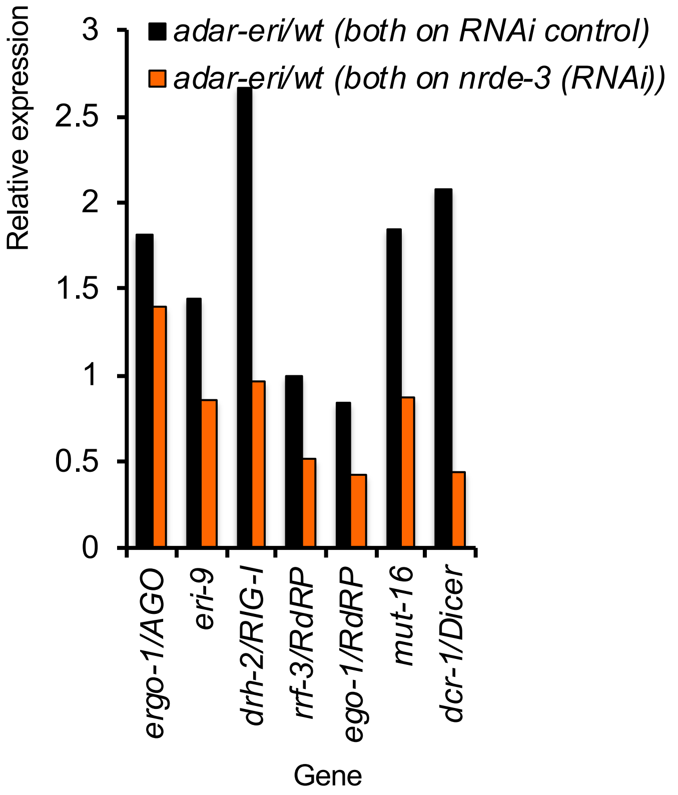

Supplemental Figure 1. A. *adar-eri* mutants display a partially penetrant rupture phenotype (blue arrows indicate rupture of internal organs through the vulva). B. *adar-eri* mutants produce novel siRNAs that do not act in canonical endogenous siRNA pathways. siRNAs are classified by the specific Argonaute protein they are known to bind to or depend on. C. *adar-eri* mutants produce more 22G small RNAs and relatively fewer 23U microRNAs. D. *eri-6/7* is required for a subset of endogenous siRNAs*.* The number of siRNAs per gene in *eri-6/7* is plotted against the number of siRNAs in wild type. E. Previously identified siRNAs that are found in the absence of *adr-1* and *adr-2* (ARL), are not further increased in *adar-eri* mutants. F. Overlap between genes producing increased or decreased numbers of siRNA in *eri-6, adar* or *adar-eri* mutants. Cutoff P<0.05. G. RNAi factor induction in *adar* and *adar-eri* mutants. The expression is relative to wild type (All P<0.01). H. Modest induction of the RNAi machinery in wild type worms exposed to control RNAi vector versus wild type worms without RNAi. I and J. RNAi factor induction in *adar-eri* triple mutants is dependent on *drh-1/RIG-I* and *nrde-3/AGO*. mRNAseq was done on wild type and *adar-eri* mutants exposed to RNAi vector control, *drh-1(RNAi)* or *nrde-3(RNAi).*

Supplemental Figure 2

2A.

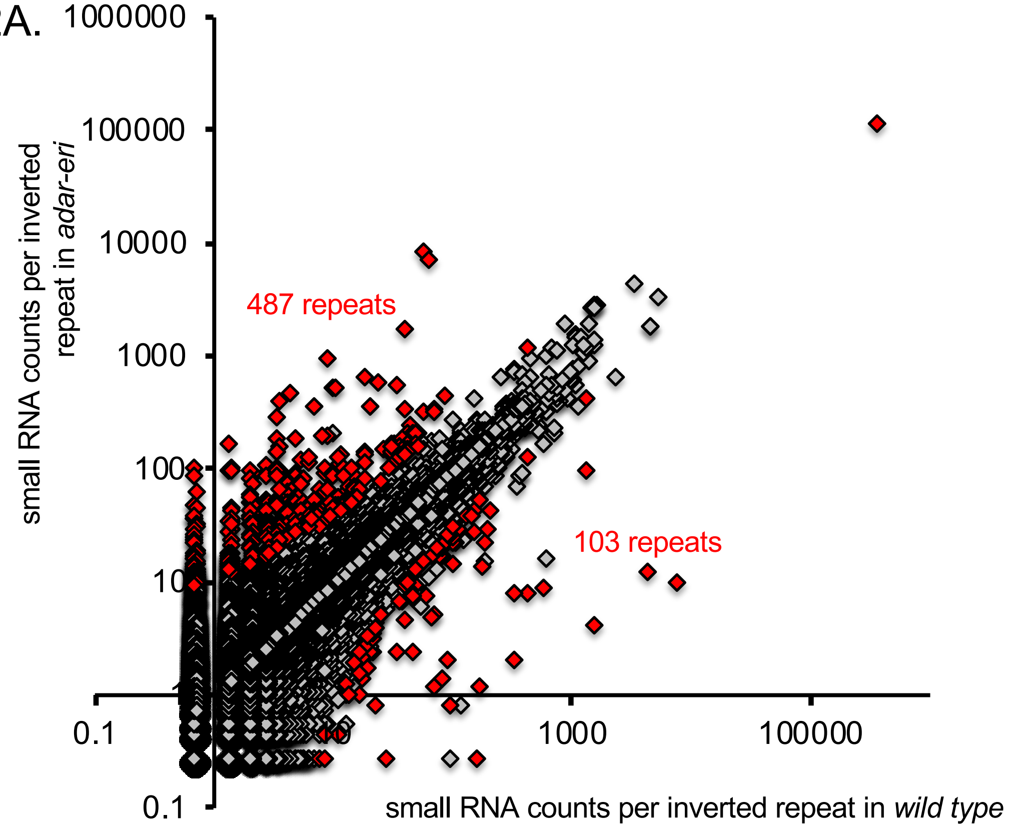

2B.

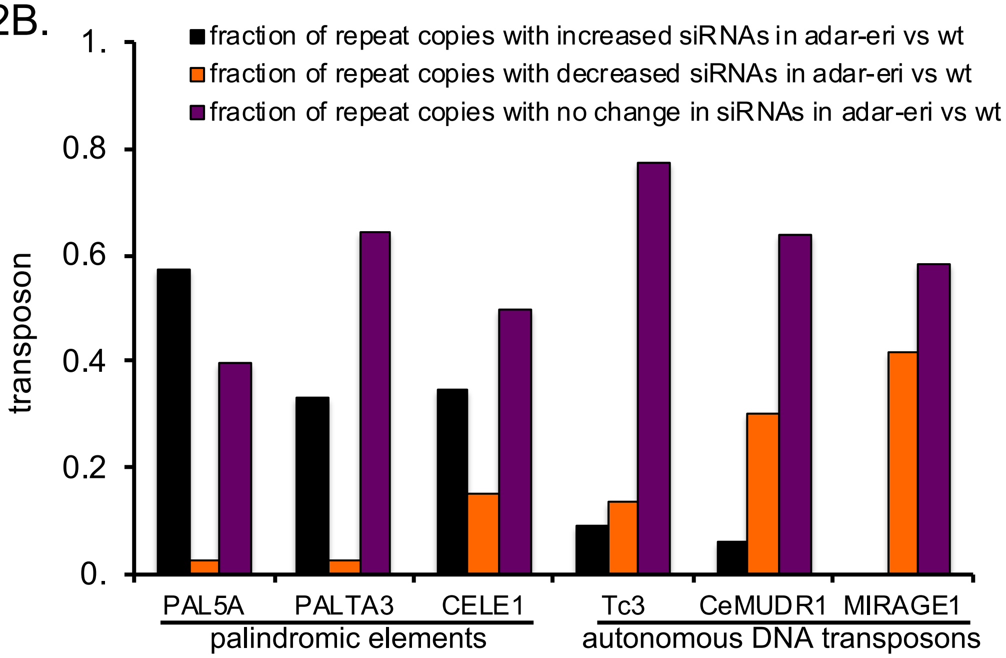

Supplemental Figure 2

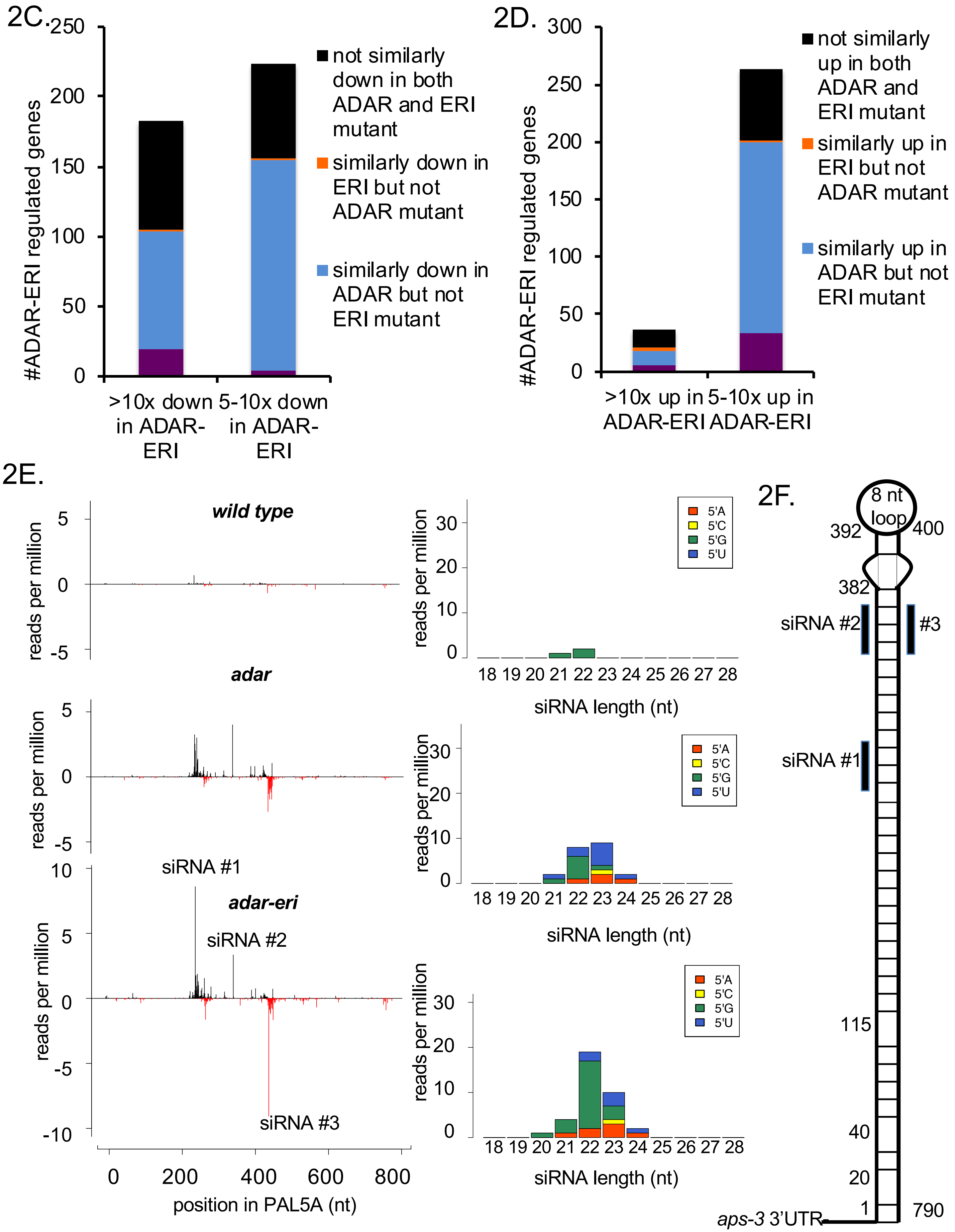

2G. CACTTAAACTTTTTCCTCACGAGGGACGAGGAAAAGGGGTTTCTAGGCCATGGCCGAGG
ATCCGTCAAGTTTCAGCGGCAATTTATCTTGCTTTGTTTTCCGCGTGTTTTCTTTTTTT
TTTCACAGATTTTCTCCCGTTTTTTCTTATCAAAACTTATAAATAAATATTTTTTGCAG
GCGCTAAAGCAATTTCCAAGTGAAAAATCGTGTATTGAGTCGGCAAGTAGCGGTGAACG
TGGGCATTGTAATATGATGGATTACGGGAATACAAAACCTAAACTTTTTCTGAAACATG
ATACATATGCTGCTTAGATGCTAAAACTACCTGATTTTCATATCGAGACCGCTGAAAAA
GTTTTAAGGTTTCCAAAATTCAACTTTTTGTTCGAAAATCTCGACTTTTTCACCAAAAA
AGTTGAATTTTGAAAATCTCAAAACTTTTTCAGCGGTCTCGTTATGAAAATCAGGTACT
TTCAGAATCTAAGCAGCATATATATCATGTTTCGGAAAAAGTTCAGGTTTGGTATTCCC
GTAATCCATCATATTACATTGCACACGTTCACCGCTACTTGCCGACTGAATACATAATT
TTTCACTCGGAAATTGCTTTAGCATCTGCAAAAAAATATTTATTCATCAGTTTTAATAA
GAAAAAACGGGGGAAAATCGGTGAAAAACACAAGAAAACACTCGGAAAAGAAAGCAAG
ATAAATTGCCGCTGAAACTTGTCGGATCCTCGGACATGGCCTAGAAACCGCATTTCCTC
GTCCCTCGTTCGAAAATAGTTGCAGTG

2H.

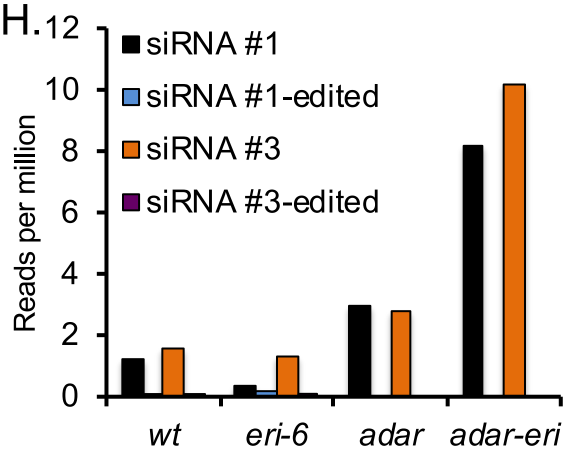

Supplemental Figure 2. A. siRNA originate from inverted repeats in *adar-eri* mutants. B. Non-autonomous, palindromic transposons (*e.g.* PAI5A, TALTA3, CELE1) produce siRNAs in *adar-eri* mutants. Autonomous DNA transposons (*e.g.* Tc3, CeMURD1 and MIRAGE1) do not produce increased siRNAs in *adar-eri* mutants. C. The extent of gene down regulation in *adar-eri, adar* and *eri* mutants. D. The extent of gene upregulation in *adar-eri, adar* and *eri* mutants. E. siRNAs mapping to the PAL5A elements in the 3'UTR of *aps-3.* sense siRNAs are black, antisense are red. The 5' nucleotide and length of siRNAs are shown. F. PAL5A folds into a hairpin structure as predicted by RNA Fold. Indicated are regions where 3 or more bases do not basepair. The 3 most abundant siRNAs in *adar-eri* are indicated as bars. mRNAseq shows this PAL5A element is part of the *aps-3* 3'UTR. G. 792 nt sequence of PAL5A in the *aps-3* 3'UTR. In bold are indicated bases not predicted to base-pair. In red are indicated 87 edited nucleotides. Editing was analyzed in mapping experiments allowing for 4 mismatches. For each edited nucleotide, editing ranges between 25% and 80%. Edits are absent in *adar* and *adar-eri* libraries. Underlined are siRNAs found most abundant in *adar-eri*. H. Abundance of the most abundant PAL5A-*aps-3* siRNAs. *wt* and *eri-6* produce fewer of these siRNAs, also when potentially edited versions of these siRNAs are included, and mapping allows for 3 mismatches.

### Supplemental Figure 3

3A.

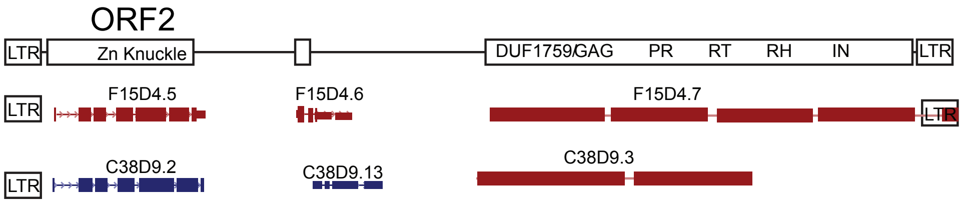

| | **Cer19-ORF2** | **Cer19-gag-pol** |
|---|---|---|
| Function | Nucleocapsid? | Gag-Pol |
| *C. elegans* | F15D4.5, C38D9.2 | F15D4.7, C38D9.3 |
| *C. brenneri* | e.g. CBN11868 | e.g. CBN0703 |
| *C. nigoni* | e.g. Cnig_chr_III.g9154 | e.g. Cnig_chr_III.g9151+others |
| *C. inopinata* | e.g. SP34_40029700 | e.g. SP34_40029600 (Gag-Pol-Env) |

| | |
|---|---|
| C.e. Cer19-ORF2(a) | **C**TF**C**GLYN**H**TSEK**C**KKFFTWFDRRERLTEMKKCHHCLE |
| C.e. Cer19-ORF2(b) | **C**AF**C**GLKN**H**TSEQ**C**RRVSTWFDRREKLTEQKKCHQCLE |
| C.n. Cer19-ORF2 | **C**AF**C**GGYS**H**QSED**C**EKKSCWRVRRMMIANKSKCFACLQ |
| C.b. Cer19-ORF2 | **C**SF**C**S-GS**H**RSYK**C**EAFGTWQERRHFLLTEKGCLKCLK |
| C.i. Cer19-ORF2 | **C**AF**C**-LKN**H**YSDN**C**GRYVTVEERKRALKKNGRCETCVA |

3B.

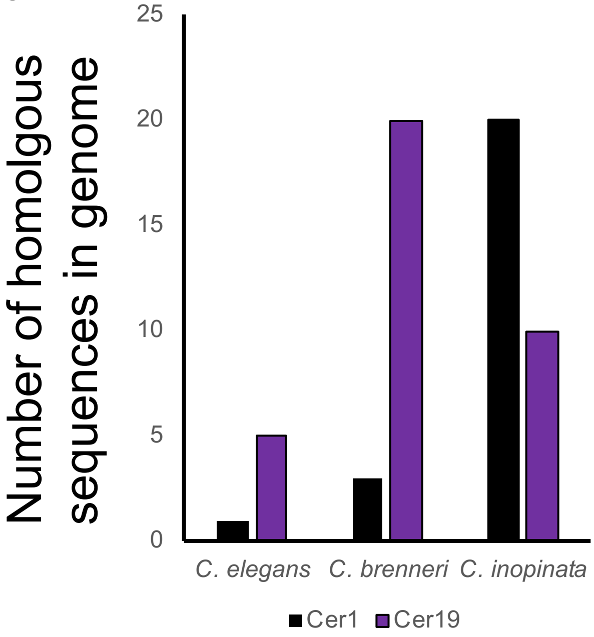

3C.

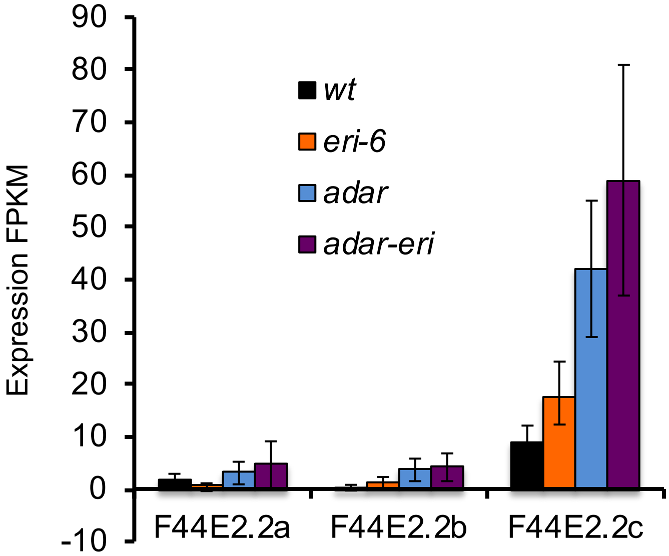

Supplemental Figure 3

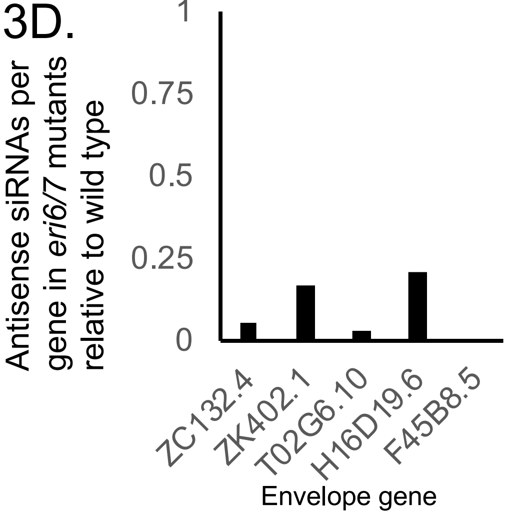

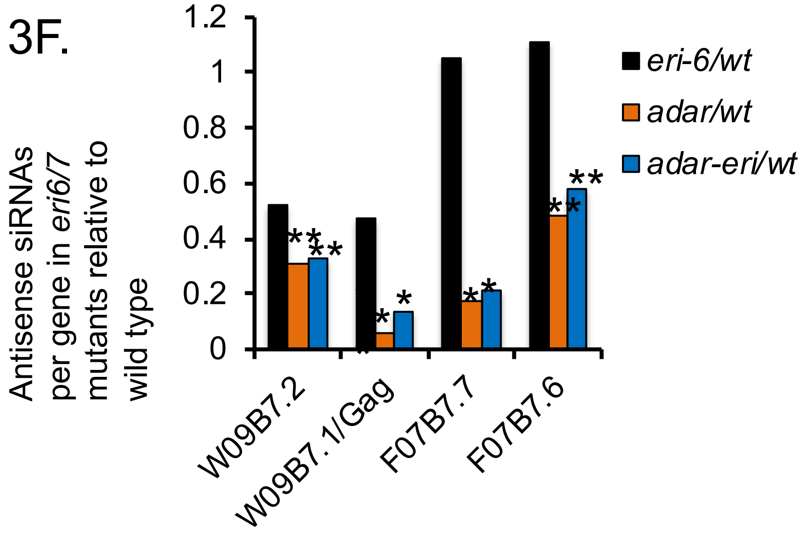

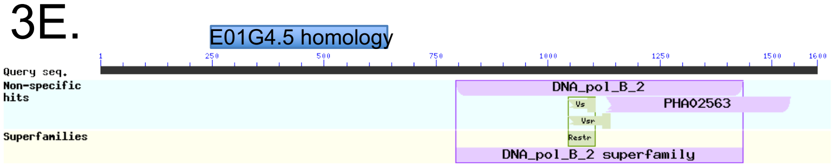

hypothetical protein B9Z55_028556 [Caenorhabditis nigoni]

| Blastp hits with B9Z55_028556 | score | number of hits |
|---|---|---|
| Caenorhabditis nigoni hits | 3369 | 36 |
| Caenorhabditis remanei hits | 1701 | 74 |
| Caenorhabditis brenneri hits | 1633 | 10 |
| Caenorhabditis elegans hits | 1528 | 2 |
| Caenorhabditis latens hits | 1469 | 10 |

Supplemental Figure 3. A. Cer19-ORFs have zf-CCHC/gag-knuckles and may encode nucleocapsid proteins. Multiple copies of Cer19 retrotransposons exist in other *Caenorhabdites C. nigoni* (*C.n.*), *C. brenneri (C.b.)* and *C. inopinata* (*C.i.*), that have a similar segmented genome containing Cer19-ORF2 and Cer19-gag-pol; the Cer19-gag-pol gene, also encodes an envelope protein in *C. inopinata*. B. Number of sequences homologous to Cer1 and Cer19 in the genomes of *C. brenneri* and *C. inopinata*. C. Absolute expression levels of Cer1 transcripts wild type, *adar, eri* and *adar-eri*. D. Envelope gene siRNAs are depleted in *eri-6/7* mutants. E. viral DNA polymerases; shown is a *C. nigoni* protein with homology to the ERI-6/7 target E01G4.5, and the number of homologs in other *Caenorhabdites*. F. siRNAs to Cer9 genes are reduced in *adar-eri* mutants (*:P<0.05, **:P<0.1).

Supplemental Figure 4

4A.

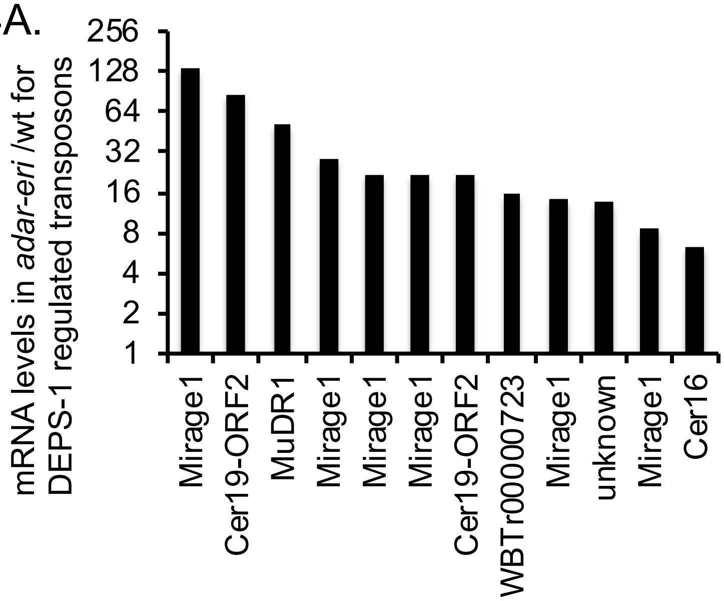

4B.

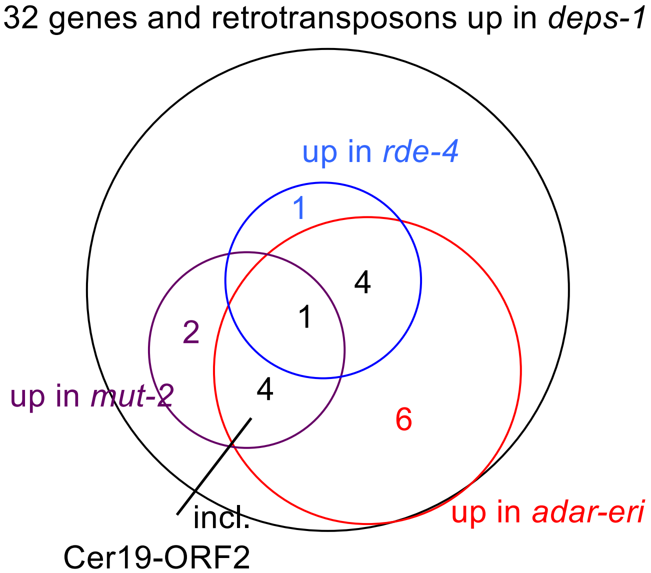

4C.

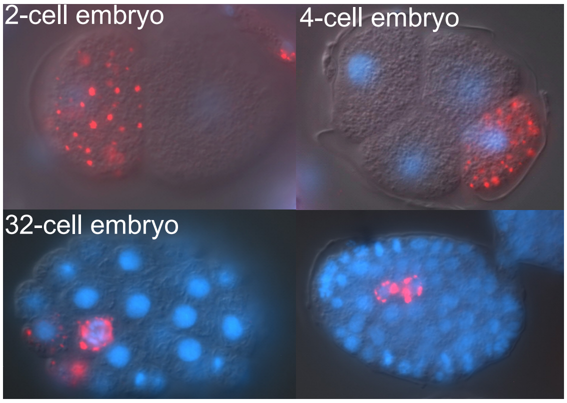

4D.

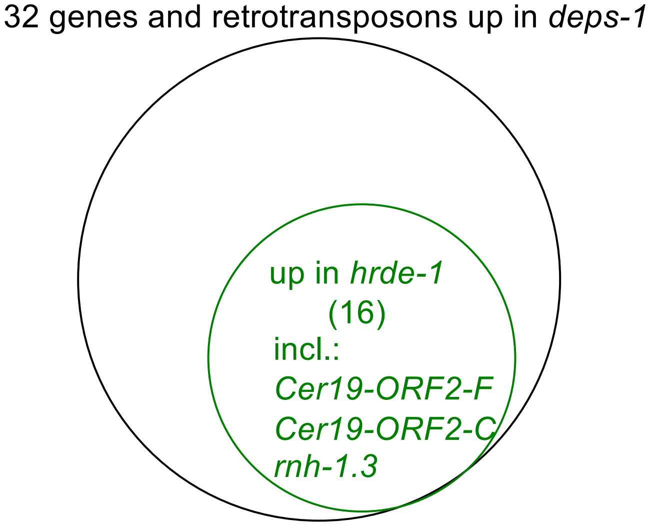

4E.

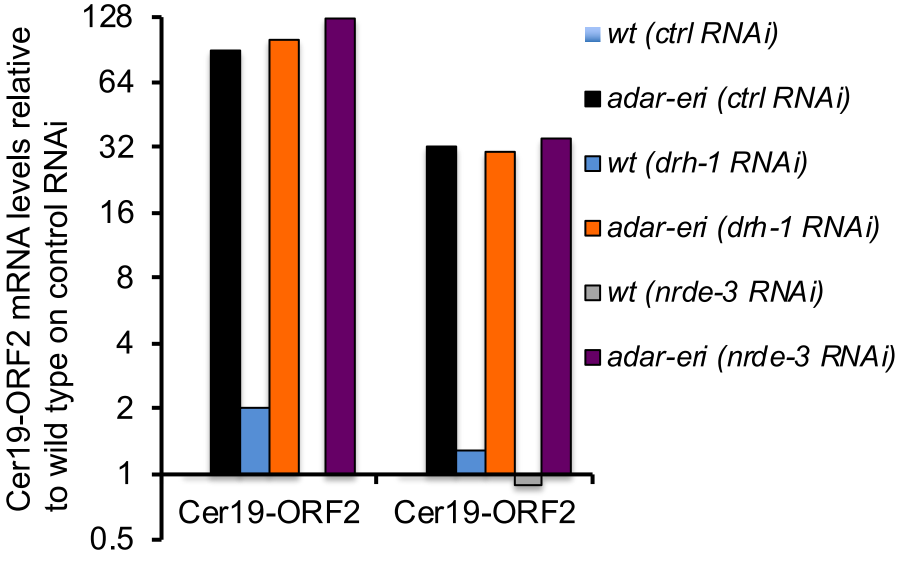

Supplemental Figure 4. P granule components and RNAi factors silence retrotransposons. A. mRNA levels are increased in *adar-eri* triple mutants versus wild type for DEPS-1-regulated retrotransposons. B. RNAi factors RDE-4 and MUT-2 regulate retrotransposons. C. PGL-1 immunohistochemistry in *adar-eri* mutant embryos (anti-PGL-1 in red, DAPI staining in blue). D. Genes silenced by *deps-1* and *hrde-1* overlap. E. *drh-1* and *nrde-3* are not required for retrotransposon silencing in wild type animals and are also not required for retrotransposon desilencing in *adar-eri* mutants.

### Supplemental Figure 5

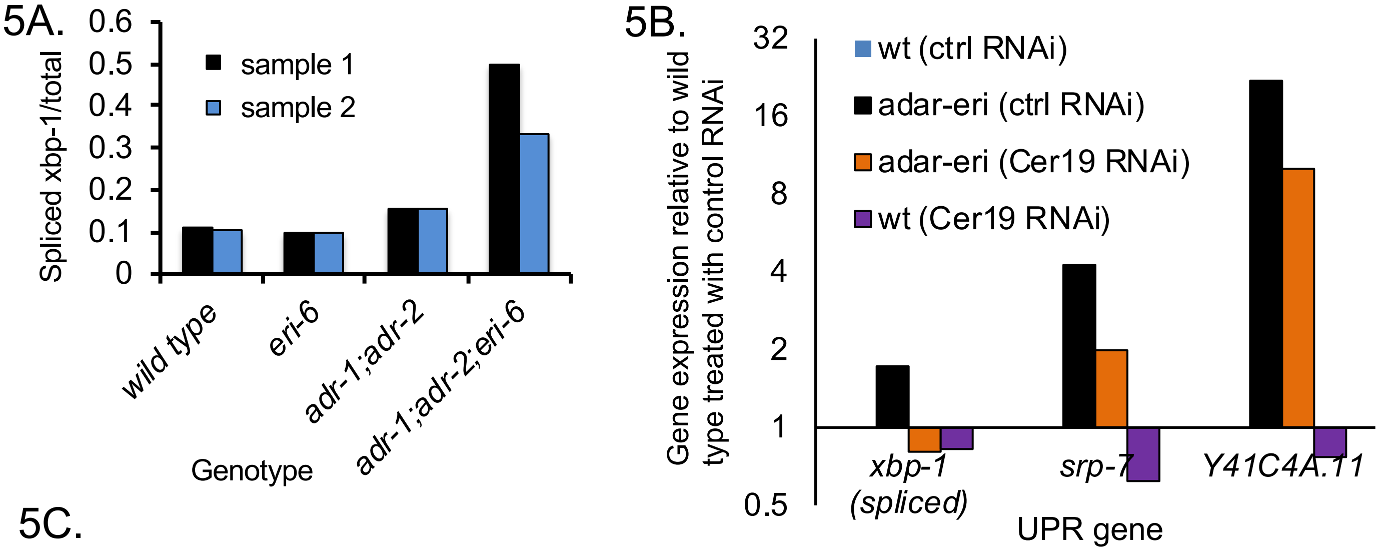

5C.

| Gene/mutation/RNAi | Function | phenotype in *adar-eri* | phenotype in *eri-6/7 (mg379)* | phenotype in *adr-1 (tm668)* |
|---|---|---|---|---|
| *hsp-4 (gk514)* | UPR | N/A | Rup, Ste at 15ºC | Rup, Ste at 15ºC |
| *hsf-1 (RNAi)* | Heat shock response | Lvl; dauer-like larvae | Ste | N/A |
| *hsf-1 (sy441)* | Heat shock response | N/A | dauer-like larvare, Lva at 22ºC, enhanced RNAi | N/A |

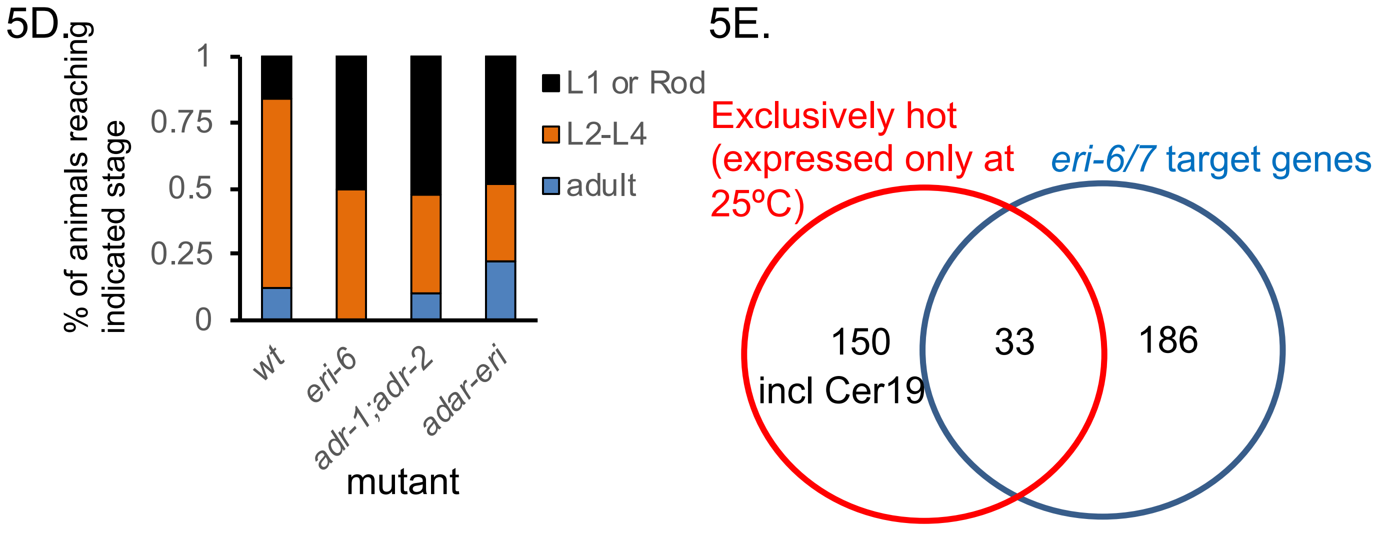

Supplemental Figure 5. A. Expression of IRE-1-spliced *xbp-1* mRNA over total *xbp-1* mRNA as measured by qRT-PCR. B. UPR gene expression in animals depleted for Cer19 RNA by RNAi, or in animals treated with control dsRNA. C. Interactions between *adar* and *eri* mutations and UPR gene depletion. Lvl: Larval lethal, Rup: rupture, Ste: sterile, Lva: Larval arrest. D. Sensitivity to 10 mg/ml tunicamycin after exposure of embryos at 25ºC. The life stages reached were scored, with L1 or Rod indicating high sensitivity to tunicamycin, while animals reaching adulthood were less sensitive. n>33. E. Cer19 and many *eri-6/7* target genes are expressed exclusively at 25ºC and not at 15 or 20ºC  (data: (32)).

### Supplemental Figure 6.

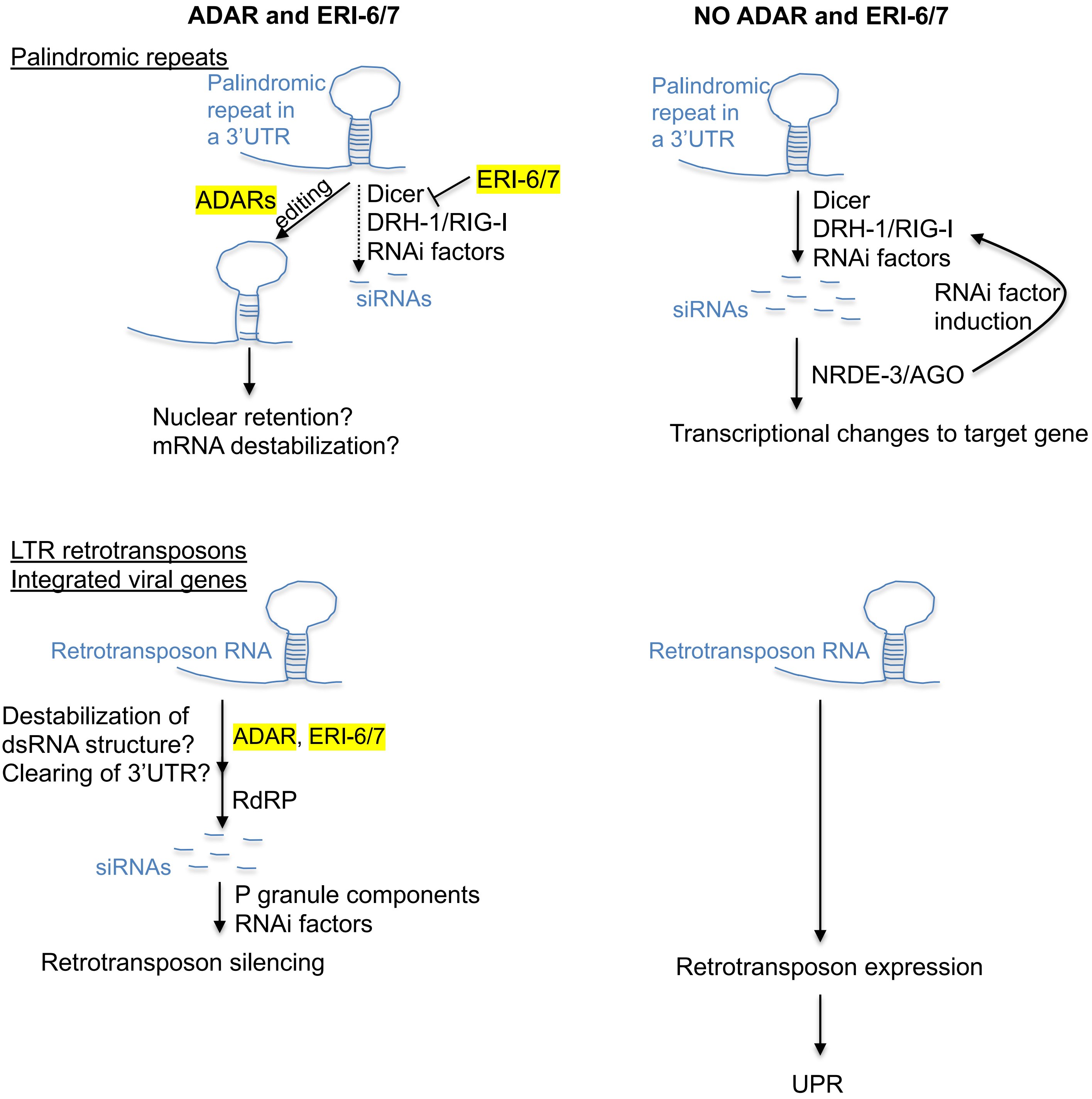

Supplemental Figure 6. Model of ADAR and ERI-6/7 activity on palindromic repeats and LTR retrotransposons.

Supplemental Table 1. The most highly mis-regulated genes in *adar-eri* triple mutants are similarly mis-regulated in *eri-6/7* and *adar* mutants

| Gene name | *wt* | *eri-6/wt* | *adar/wt* | *adar-eri/wt* |
|---|---|---|---|---|
| *C17E7.9* | | **14** | **18** | **29** |
| *C17E7.13* | | **14** | **18** | **29** |
| *T05E12.10* | | **23** | **5** | **25** |
| *F19B10.10* | | **6** | **4** | **23** |
| *Y39G8B.2* | | **12** | **3** | **21** |
| *T02G6.5* | | **6** | **20** | **18** |
| *W02B12.12* | | **11** | **12** | **17** |
| *Y47D3A.13* | | **9** | **11** | **17** |
| *F48C1.9* | | **37** | **7** | **15** |
| *K08C9.2* | | **9** | **11** | **15** |
| *F26B1.8* | | **11** | **11** | **15** |
| *spe-11* | | **10** | **12** | **14** |
| *drh-2* | | **3** | **11** | **14** |
| *T10E9.4* | | **9** | **11** | **14** |
| *F02H6.3* | | **10** | **19** | **14** |
| *C18D4.6* | | **12** | **9** | **13** |
| *F46C3.4* | | **7** | **15** | **13** |
| *C10G11.8* | | **10** | **12** | **12** |
| *F28H6.4* | | **4** | **9** | **12** |
| *K09E3.5* | | **7** | **14** | **12** |
| *nspd-1* | | **15** | **10** | **12** |

fold change over wild type

| 1.00 | 4 | 10 |
|---|---|---|

| Gene name | *wt* | *eri-6/wt* | *adar/wt* | *adar-eri/wt* |
|---|---|---|---|---|
| *F59F5.4* | | | | |
| *F54D5.20* | | | | |
| *T22B7.13* | | | | |
| *C25F9.17* | | | | |
| *H14A12.9* | | | | |
| *C17C3.24* | | | | |
| *ins-21* | | | | |
| *gst-33* | | | | |
| *F08D12.2* | | | | |
| *cyp-35A4* | | | | |
| *T19H12.6* | | | | |
| *mtl-2* | | | | |
| *clec-206* | | | | |
| *Y110A2AL.9* | | | | |
| *cyp-35A3* | | | | |

fold change over wild type

| 1 | 0.10 | 0.01 |
|---|---|---|

Supplemental Table 2. Expression of Cer19 and Cer1 retrotransposons in of *adar, eri-6/7* and *adar-eri* mutants.

| gene_short_name | wt_FPKM | eri-6_FPKM | adr1adr2_FPKM | adr1adr2eri6_FPKM |
|---|---|---|---|---|
| C38D9.2 | 0.25 | 0.34 | 4.99 | 21.92 |
| F15D4.5 | 0.95 | 0.79 | 10.50 | 20.37 |
| Cer1 | 11.15 | 19.17 | 48.74 | 69.46 |

### Supplemental Table 3. Loci that produce the highest numbers of siRNAs (RPM)

| WB gene name | average wt (RPM) | average eri6 (RPM) | eri6/wt | pvalue | gene name | seq name | |
|---|---|---|---|---|---|---|---|
| WBGene00219699 | 85213.00857 | 1286.016492 | 0.01509179 | 3.2939E-11 | linc-22 | F47E1.18 | eri target |
| WBGene00003332 | 56540.22193 | 13363.45277 | 0.23635303 | 0.10470101 | mir-239.1 | C34E11.5 | eri target |
| WBGene00189951 | 46943.47544 | 44815.53686 | 0.9546702 | 0.94378338 | linc-29 | Y73B6BL.269 | |
| WBGene00194660 | 45236.61856 | 37285.68892 | 0.82423687 | 0.77818466 | 0 | W09C5.12 | |
| WBGene00009511 | 42010.20104 | 31727.62346 | 0.75523617 | 0.66858616 | 0 | F37H8.2 | |
| WBGene00014884 | 40586.41236 | 45749.24919 | 1.12720604 | 0.85587885 | 0 | Y39E4B.14 | |
| WBGene00013228 | 34564.78615 | 20261.85194 | 0.58619926 | 0.44357247 | 0 | Y56A3A.7 | |
| WBGene00196028 | 33587.16392 | 19364.763 | 0.57655249 | 0.47960294 | 0 | C32D5.15 | |
| WBGene00006771 | 33257.25575 | 31113.09439 | 0.93552801 | 0.91723849 | tln-1 | Y71G12B.11 | |
| WBGene00000801 | 32474.0039 | 30860.61543 | 0.95031754 | 0.94088584 | cars-2 | Y23H5A.1 | |
| WBGene00199017 | 31876.40943 | 25858.10721 | 0.81119887 | 0.75578258 | 0 | Y71G12B.36 | |
| WBGene00009247 | 29190.3242 | 29124.66386 | 0.99775061 | 0.99725069 | bath-45 | F29C12.5 | |
| WBGene00020146 | 29062.1836 | 31903.51144 | 1.09776718 | 0.89616545 | got-1.2 | T01C8.5 | |
| WBGene00009509 | 27111.58896 | 25769.01607 | 0.95047974 | 0.94234628 | 0 | F37D6.3 | |
| WBGene00001433 | 26937.75604 | 23202.55535 | 0.86133958 | 0.8240591 | fkb-8 | Y18D10A.25 | |
| WBGene00015012 | 26363.3494 | 23517.36555 | 0.89204771 | 0.85697115 | bath-20 | B0047.1 | |
| WBGene00198071 | 26157.01323 | 13978.2295 | 0.534397 | 0.42997735 | 0 | C12C8.6 | |
| WBGene00010475 | 25615.45618 | 31257.53185 | 1.2202606 | 0.78005441 | 0 | K01F9.2 | |
| WBGene00199557 | 25597.50457 | 31232.85631 | 1.22015239 | 0.78023152 | 0 | K01F9.7 | |
| WBGene00014836 | 25485.54043 | 29074.24303 | 1.14081328 | 0.84053903 | 0 | T09F5.12 | |
| WBGene00012228 | 25394.69643 | 23855.26071 | 0.93937964 | 0.92764348 | 0 | W03H9.2 | |
| **WBGene00008862** | **25233.52826** | **21444.29083** | **0.84983323** | **0.803713** | **0** | **F15D4.5** | **Cer19-ORF2** |
| WBGene00007125 | 25152.49751 | 26691.66264 | 1.06119333 | 0.92686474 | 0 | B0250.8 | |
| WBGene00019517 | 24940.39287 | 25536.37457 | 1.02389624 | 0.97282105 | 0 | K08B4.5 | |
| WBGene00011737 | 22248.01701 | 23257.73881 | 1.0453848 | 0.94760589 | sqst-1 | T12G3.1 | |
| WBGene00195155 | 21691.71599 | 26744.46383 | 1.23293445 | 0.74613101 | 0 | W04A8.9 | |
| WBGene00009310 | 21654.36073 | 19853.0367 | 0.91681472 | 0.9040819 | 0 | F32A11.5 | |
| WBGene00021606 | 21201.6122 | 16093.0618 | 0.75904897 | 0.68417912 | 0 | Y46H3C.6 | |
| WBGene00206497 | 20145.48142 | 16836.30123 | 0.83573586 | 0.77428718 | 0 | Y52B11A.18 | |
| WBGene00219589 | 19988.53018 | 14356.38729 | 0.71823126 | 0.60935123 | anr-35 | F37H8.13 | |
| WBGene00004622 | 19835.06154 | 19433.92164 | 0.97977622 | 0.97619643 | rrn-3.1 | F31C3.9 | |
| WBGene00219207 | 18750.30694 | 20614.72695 | 1.09943411 | 0.8827211 | 0 | C01F6.14 | |
| WBGene00022163 | 18521.55627 | 20615.77923 | 1.11306949 | 0.87487087 | 0 | Y71G12B.28 | |
| WBGene00206415 | 18192.63094 | 17384.55715 | 0.95558236 | 0.94516441 | 0 | F59H6.14 | |
| WBGene00044161 | 17732.47001 | 16442.08273 | 0.92723026 | 0.90851073 | 0 | C47G2.8 | |
| WBGene00077437 | 17450.49452 | 1902.877761 | 0.10904435 | 0.00045635 | 0 | Y59H11AR.6 | eri target |
| WBGene00011964 | 17343.44452 | 18103.68157 | 1.04383426 | 0.94705232 | saeg-2 | T23G5.6 | |
| WBGene00021673 | 16755.26055 | 15157.28374 | 0.90462835 | 0.8800504 | 0 | Y48G1BM.8 | |
| WBGene00014663 | 16610.2247 | 18815.37945 | 1.13275888 | 0.8564527 | 0 | C01G6.10 | |
| WBGene00013685 | 16583.34258 | 16253.89586 | 0.98013388 | 0.9765931 | 0 | Y105E8A.28 | |
| WBGene00012578 | 16519.6882 | 12758.91073 | 0.77234574 | 0.71715247 | 0 | Y37H9A.3 | |
| WBGene00021098 | 16316.83292 | 16895.87622 | 1.03548748 | 0.95874193 | fbxb-14 | W08F4.9 | |
| WBGene00000031 | 16080.14483 | 13012.11676 | 0.80920395 | 0.7251801 | abu-8 | C03A7.14 | |
| WBGene00012577 | 15575.72221 | 11926.90498 | 0.76573688 | 0.70925415 | 0 | Y37H9A.2 | |
| WBGene00001976 | 15465.53348 | 16547.70511 | 1.06997312 | 0.9213362 | hmg-11 | T05A7.4 | |
| WBGene00044907 | 14798.7007 | 14485.23092 | 0.97881775 | 0.97414505 | 0 | K08E7.10 | |
| WBGene00012961 | 14579.92355 | 13282.88566 | 0.91103946 | 0.8779299 | 0 | Y47H10A.5 | |
| WBGene00007309 | 14413.50258 | 8135.5162 | 0.56443714 | 0.35396457 | 0 | C04G2.10 | |
| WBGene00045366 | 14038.59573 | 746.1940019 | 0.05315304 | 0.00021296 | 0 | Y17D7C.3 | eri target |
| WBGene00013221 | 13933.2449 | 14086.15113 | 1.0109742 | 0.9884593 | 0 | Y54G11A.14 | |
| WBGene00019735 | 13775.17415 | 13888.09358 | 1.00819731 | 0.9905849 | 0 | M02F4.1 | |
| WBGene00044670 | 13701.28392 | 14100.16974 | 1.02911303 | 0.96193018 | rac-3 | K03D3.13 | |
| WBGene00021736 | 13291.80748 | 14038.97489 | 1.05621263 | 0.93172662 | wrb-1 | Y50D4A.2 | |
| WBGene00004041 | 13165.16446 | 9981.258433 | 0.75815676 | 0.71066674 | plg-1 | F44E2.11 | |
| WBGene00001898 | 13150.04033 | 13033.32846 | 0.9911246 | 0.98876326 | his-24 | M163.3 | |
| WBGene00013152 | 12987.11273 | 10579.77937 | 0.81463676 | 0.7567054 | 0 | Y53F4B.5 | |
| WBGene00012040 | 12795.86174 | 12144.42652 | 0.94909016 | 0.94016356 | 0 | T26E3.7 | |
| WBGene00001859 | 12687.43302 | 4898.602561 | 0.38609879 | 0.14098341 | hil-8 | T05E8.2 | |
| WBGene00077762 | 12639.82697 | 9897.562562 | 0.78304573 | 0.7082479 | 0 | Y57G11C.499 | |
| WBGene00018648 | 12572.21038 | 11800.8561 | 0.93864609 | 0.91822057 | 0 | F49F1.8 | |
| WBGene00255736 | 12459.08101 | 12909.56862 | 1.03615737 | 0.95817194 | 0 | C03H5.10 | |
| WBGene00007376 | 12439.22677 | 12072.79695 | 0.9705424 | 0.96337809 | 0 | C06C3.5 | |
| WBGene00019541 | 12367.99248 | 12049.20189 | 0.97422455 | 0.96915862 | 0 | K08F11.1 | |
| WBGene00009788 | 12356.03855 | 11939.35042 | 0.96627656 | 0.96029948 | 0 | F46F2.4 | |
| WBGene00016635 | 12199.67578 | 244.1324066 | 0.02001138 | 2.7273E-10 | 0 | C44B11.6 | eri target |
| WBGene00043534 | 12138.03033 | 8230.706524 | 0.67809243 | 0.54408493 | 0 | W02B12.13 | |
| WBGene00219725 | 11957.21237 | 13003.85882 | 1.08753265 | 0.89553649 | linc-63 | T03F1.13 | |
| WBGene00014914 | 11956.3872 | 12242.12788 | 1.02389858 | 0.97168251 | 0 | Y52B11A.11 | |
| WBGene00219707 | 11843.34933 | 327.8854234 | 0.02768519 | 1.3616E-08 | linc-8 | F59H5.6 | eri target |
| WBGene00008517 | 11611.32693 | 12864.50431 | 1.10792715 | 0.88098517 | 0 | F02C12.3 | |

Supplemental Table 4. Viral envelope proteins in *C. inopinata* homologous to the protein encoded by ZC132.4.

| C. inopinata gene | Score (bits) | E value |
|---|---|---|
| Sp34_30057800 | 472 | e-133 |
| Sp34_X0163400 | 356 | 3e-98 |
| Sp34_20074700 | 354 | 7e-98 |
| Sp34_X0162300 | 353 | 2e-97 |
| Sp34_X0151100 | 353 | 2e-97 |
| Sp34_10027800 | 353 | 2e-97 |
| Sp34_50062500 | 341 | 6e-94 |
| Sp34_30153900 | 337 | 1e-92 |
| Sp34_X0048100 | 336 | 3e-92 |
| Sp34_40227400 | 332 | 6e-91 |
| Sp34_10203300 | 325 | 7e-89 |
| Sp34_30042200 | 323 | 2e-88 |
| Sp34_40029600 | 311 | 9e-85 |
| Sp34_40163100 | 310 | 2e-84 |
| Sp34_40305100 | 309 | 3e-84 |
| Sp34_10208800 | 308 | 5e-84 |
| Sp34_20180500 | 307 | 2e-83 |
| Sp34_30116500 | 306 | 2e-83 |
| Sp34_40001300 | 301 | 1e-81 |
| Sp34_40002400 | 300 | 1e-81 |
| Sp34_10245910 | 296 | 3e-80 |
| Sp34_X0122100 | 295 | 5e-80 |
| Sp34_20155900 | 295 | 5e-80 |
| Sp34_10207710 | 293 | 2e-79 |
| Sp34_30095600 | 293 | 2e-79 |
| Sp34_20086700 | 292 | 5e-79 |
| Sp34_10205300 | 278 | 9e-75 |
| Sp34_20198700 | 276 | 4e-74 |
| Sp34_10195000 | 229 | 4e-60 |
| Sp34_10130300 | 228 | 6e-60 |
| Sp34_10193200 | 223 | 2e-58 |
| Sp34_10204100 | 221 | 7e-58 |
| Sp34_30159700 | 211 | 1e-54 |
| Sp34_20202800 | 202 | 5e-52 |
| Sp34_30153500 | 177 | 3e-44 |
| Sp34_50417510 | 174 | 1e-43 |
| Sp34_30191300 | 168 | 1e-41 |
| Sp34_X0176200 | 166 | 4e-41 |
| Sp34_X0243410 | 154 | 1e-37 |
| Sp34_30138000 | 153 | 4e-37 |
| Sp34_10163600 | 145 | 7e-35 |
| Sp34_10143800 | 140 | 2e-33 |
| Sp34_50397600 | 136 | 4e-32 |
| Sp34_X0181200 | 128 | 1e-29 |
| Sp34_X0146100 | 128 | 1e-29 |
| Sp34_10193800 | 125 | 6e-29 |
| Sp34_10117000 | 125 | 1e-28 |
| Sp34_10194900 | 124 | 2e-28 |
| Sp34_10208200 | 120 | 2e-27 |
| Sp34_20059000 | 120 | 3e-27 |
| Sp34_20171500 | 117 | 2e-26 |
| Sp34_10206200 | 114 | 2e-25 |
| Sp34_10195900 | 113 | 4e-25 |
| Sp34_50207100 | 110 | 3e-24 |
| Sp34_30102400 | 105 | 8e-23 |
| Sp34_50088100 | 94 | 2e-19 |
| Sp34_30176800 | 91 | 1e-18 |
| Sp34_30236000 | 80 | 4e-15 |

Supplemental Table 5. Genes induced upon Orsay virus infection that are co-regulated with retrotransposons.

| WB Gene name | Sequence Gene name | Gene name |
|---|---|---|
| WBGene00007660 | C17H1.7 | pals-6 |
| WBGene00045401 | T26F2.3 | eol-1 |
| WBGene00015227 | B0507.10 | ddn-1 |
| WBGene00017214 | F07E5.9 | 0 |
| WBGene00015225 | B0507.8 | 0 |
| WBGene00012399 | Y6E2A.5 | 0 |
| WBGene00021081 | W08A12.4 | pals-33 |
| WBGene00008267 | C53A5.9 | 0 |
| WBGene00018345 | F42C5.3 | 0 |
| WBGene00045393 | F26D11.13 | 0 |
| WBGene00007132 | B0284.2 | pals-27 |
| WBGene00138721 | C54D10.14 | pals-37 |
| WBGene00018353 | F42G2.4 | fbxa-182 |
| WBGene00194713 | F19B10.13 | 0 |
| WBGene00015602 | C08E3.10 | fbxa-158 |
| WBGene00008302 | C54D10.8 | pals-38 |
| WBGene00009835 | F47H4.2 | 0 |
| WBGene00009908 | F49H6.5 | 0 |
| WBGene00004509 | M01G12.12 | rrf-2 |
| WBGene00022674 | ZK177.9 | 0 |
| WBGene00007694 | C23H4.6 | 0 |
| WBGene00017705 | F22E5.6 | 0 |
| WBGene00022570 | ZC239.12 | sdz-35 |
| WBGene00011052 | R06B9.1 | arrd-11 |
| WBGene00001577 | T12A7.1 | gem-4 |
| WBGene00021654 | Y47G7B.2 | 0 |
| WBGene00202499 | B0507.15 | 0 |
| WBGene00008726 | F13A7.11 | 0 |
| WBGene00004811 | F47H4.10 | skr-5 |
| WBGene00008850 | F15B9.6 | 0 |
| WBGene00000841 | K08E7.7 | cul-6 |
| WBGene00010752 | K10G4.3 | 0 |
| WBGene00009957 | F53B2.8 | 0 |
| WBGene00007440 | C08E8.4 | 0 |
| WBGene00000503 | C04F6.3 | cht-1 |
| WBGene00016788 | C49G7.10 | 0 |
| WBGene00007133 | B0284.3 | 0 |
| WBGene00006987 | EGAP1.3 | zmp-1 |
| WBGene00015537 | C06E4.8 | 0 |
